# A low-annotation-budget PubMedBERT classifier for chondrogenesis regulator discovery via active learning

**DOI:** 10.64898/2026.09.24.754045

**Authors:** Ana-Mariya Anhel, Mandy Jayne Peffers, David A Young, Jamie Soul

## Abstract

**Motivation:** Biomedical natural language processing (Bio-NLP) classification tasks are often limited by the cost of manual annotation, especially for specialised extraction problems where labelled corpora are scarce. Active learning can reduce this cost by prioritising informative samples, while large language models (LLMs) avoid task-specific annotation, but at high computational cost. We ask whether an encoder trained via active learning can match LLM performance at a lower computational cost, using identification of chondrogenesis regulators from PubMed abstracts as a test case applicable to other under-annotated Bio-NLP problems.

**Results:** We built an active learning framework to fine-tune PubMedBERT to classify gene mentions as regulators of chondrocyte differentiation, achieving an AUC-ROC of 0.93 using 1,238 annotated sentences. Against open-weight LLMs (Qwen3, Llama-3.1), PubMedBERT matched the top-performing LLMs on AUC-ROC and achieved the highest precision (0.69) of any model, while classifying the full corpus far faster. Applied to 67,111 gene mentions, it identified 1,128 candidate regulators, recovered 79% of Gene Ontology-annotated genes related to chondrocyte differentiation (94 genes recovered), and proposed 981 new candidates. The pipeline is agnostic to the target process, suggesting it could be adapted to identify regulators of other biological processes.

**Availability and Implementation:** Code is available at https://github.com/ChondroTextomics/ALRegulatorDiscovery. Data are available at Zenodo: https://doi.org/10.5281/zenodo.22746090. Final model is available in HuggingFace Hub amav/pubmedbert-chondrogenesis-classifier

**Supplementary information:** Supplementary data are available in a separate file

## Introduction

Identifying genes that regulate biological processes is a key objective in functional genomics (Przybyla and Gilbert 2022). Machine learning methods are often used to prioritise candidate genes (Moreau and Tranchevent 2012, Balachandran *et al*. 2024, Yu *et al*. 2024, Schipper *et al*. 2025, Wu *et al*. 2025). Supervised machine learning methods rely on prior knowledge, meaning they typically require a set of genes already known to be involved in the process to train, supervise, or benchmark the model (Zhao XM *et al*. 2008, Le 2020, Azadifar and Ahmadi 2022). In Gene Ontology (GO), the availability of this annotation varies across biological processes. In humans, muscle tissue development (GO:0060537) is annotated with 453 genes, whereas bone marrow development (GO:0048539) is annotated with only nine. Sparse annotation leaves too few positive examples to train supervised methods in order to predict novel regulators. Much of this missing knowledge already exists in the primary literature, suggesting that automatically extracting relationships between genes and biological processes from published texts could help expand coverage (Xing *et al*. 2018, Collins *et al*. 2024, Rai *et al*. 2024).

The extraction of genes linked to biological processes has been less explored in biomedical natural language processing (Bio-NLP) than gene-disease (Rahit *et al*. 2024, Jumaa *et al*. 2025, Collins *et al*. 2026) or protein-protein relationship extraction (Koyabu, Phan, and Ohkawa 2015, Nezamuldeen and Jafri 2023, Mehryary *et al*. 2024). Tools such as RelEX (Fundel, Ku ffner, and Zimmer 2007), BELMiner (Ravikumar, Rastegar-Mojarad, and Liu 2017), and SemRep (Kilicoglu *et al*. 2020) mainly rely on manually crafted rules, making them difficult to adapt to different biological domains (Ben Abacha and Zweigenbaum 2011, Bose *et al*. 2021).

Transformer-based language models can overcome these limitations by learning contextual representations from large corpora. Pre-training on biomedical text can further reduce the labelled data required downstream. For instance, BioBERT has achieved strong relationship-extraction performance under a constrained annotation budget (Giles *et al*. 2020). Subsequent models such as SciBERT (Beltagy, Lo, and Cohan 2019), BioELECTRA (Kanakarajan, Kundumani, and Sankarasubbu 2021), PubMedBERT (Gu *et al*. 2021) and BioLinkBERT (Yasunaga, Leskovec, and Liang 2022) have reported strong results across biomedical NLP benchmarks.

Genes associated with specific biological processes can be extracted by fine-tuning a domain encoder using labelled examples. Publicly available annotated corpora for gene-GO biological processes do exist, such as BC4GO (Van Auken *et al*. 2014) and CRAFT (Bada *et al*. 2012); however, they do not span the entire Gene Ontology terminology, so many biological processes are not represented. For many processes, the number of annotated instances is insufficient to train a classifier for that process. PubMed provides underlying evidence, but no corresponding annotations. Untargeted labelling is inefficient, as biomedical abstracts typically mention many genes, only a small proportion of which are regulators of a given process of interest. An annotation approach is therefore needed that prioritises sample selection to help achieve high model performance while minimising the amount of labelled data required.

To address this challenge, we can use active learning, a machine learning framework in which the most informative samples are iteratively selected for manual annotation and the model is retrained with them until a stopping criterion is met (Cohn, Ghahramani, and Jordan 1996), for example, when the model reaches a desired performance. This targeted selection strategy has been demonstrated across several tasks to reach comparable or better performance than random sampling for a given annotation budget (Sener and Savarese 2018, Ein-Dor *et al*. 2020, Shi, Li, and Zhou 2023, Doucet *et al*. 2025), including on biomedical NLP tasks with limited data such as entity recognition (Shi, Li, and Zhou 2023), information extraction from non-English text (S uvalov, Laur, and Kolde 2023), clinical and biomedical sequence tagging (Shelmanov *et al*. 2019), and the retrieval of functionally important protein residues from text (Vollmar *et al*. 2024). A commonly used selection strategy is hybrid sampling, which combines uncertainty and diversity sampling (He *et al*. 2014, Ash *et al*. 2019, Munro 2021, Doucet *et al*. 2025).

Positive examples, defined as genes annotated as regulators of the biological process of interest, are relatively scarce. This introduces class imbalance, which is common in biomedical datasets (Blanchard *et al*. 2022, De Angeli *et al*. 2022, Yadav *et al*. 2022), and can lead standard selection strategies to under-sample the minority class. The iterative nature of the active learning framework allows the adaptive implementation of strategies to address this imbalance (Tomanek and Hahn 2009), such as class-aware sampling, and the dynamic removal of such strategies if they are suspected of affecting the model’s performance.

LLMs provide an alternative to fine-tuning encoders that requires no task-specific annotation (Agrawal *et al*. 2022, Luo *et al*. 2022, Labrak, Rouvier, and Dufour 2024) and open-weight models reduce reliance on commercial APIs (Touvron *et al*. 2023, Bumgardner *et al*. 2024). However, computational costs remain, including memory requirements, which scale with model size. Therefore, the use of language models depends on the computational resources (processing time, hardware settings, money, etc) available to research groups. In this work, we compare the two approaches, fine-tuning an encoder using an active learning framework and using an LLM directly, in a single-GPU local-inference setting representative of environments without dedicated computing infrastructure.

We apply both to the extraction of regulators of chondrogenesis, the process by which mesenchymal stem cells (MSCs) differentiate into chondrocytes, the cells that form cartilage (Cheung *et al*. 2020, Chen M *et al*. 2024). Structured annotation of this process is limited. The GO term for chondrocyte differentiation (GO:0002062) is annotated with only 119 human genes. Processes with sparse structured coverage are precisely those for which literature-based extraction of gene–process relations are of practical value, making chondrogenesis an appropriate test case. Understanding how this process is regulated is clinically important, as it can inform strategies to enhance cartilage regeneration and lead to treatments for joint diseases such as osteoarthritis, where cartilage degeneration is a hallmark (Boeuf and Richter 2010, Welsh and Sikder 2025).

We make the following contributions to the extraction of genes related to biological processes under constrained settings: **(i)** We fine-tune PubMedBERT within an active learning loop to classify gene mentions in PubMed sentences as regulators of a target biological process, reaching robust performance from 1,238 expert-labelled sentences. **(ii)** We compare open-weight LLMs (Qwen and Llama) to a fine-tuned encoder (PubMedBERT) on the same task, reporting both classification performance and corpus-scale prediction time in a single-GPU local inference setting. **(iii)** We apply the pipeline to chondrogenesis-related PubMed abstracts, recovering known regulators and candidate genes absent from the GO annotation for cartilage development, and we release the annotated corpus and code to support extension to other sparsely annotated biological processes.

## Methods

### Data Extraction Pipeline

A multi-step data extraction pipeline, the overview of which is described in Figure 1, was implemented to retrieve sentences containing at least one recognised gene and a keyword related to chondrogenesis from a PubMed query. Relevant publications’ titles and abstracts were retrieved from PubMed using the NCBI’s E-utilities API (ESearch). Next, SciSpacy (Neumann *et al*. 2019) splitter (version 0.6.2), a pre-trained model for biomedical text. These sentences were used as input for GNorm2 (Wei *et al*. 2023), a biomedical Named Entity Recognition (NER) tool that extracts genes and links them to standardised identifiers such as NCBI Gene IDs. GNorm2 was run via the Docker image nadarajan07/gnorm2-ncbi:latest (https://hub.docker.com/r/nadarajan07/gnorm2-ncbi). Finally, only sentences that included at least one gene mention and a keyword related to chondrogenesis (for example, “development” or “differentiation”) were retained. The full list of words used to filter the sentences can be found in Supplementary Table S1.

**Figure 1.**
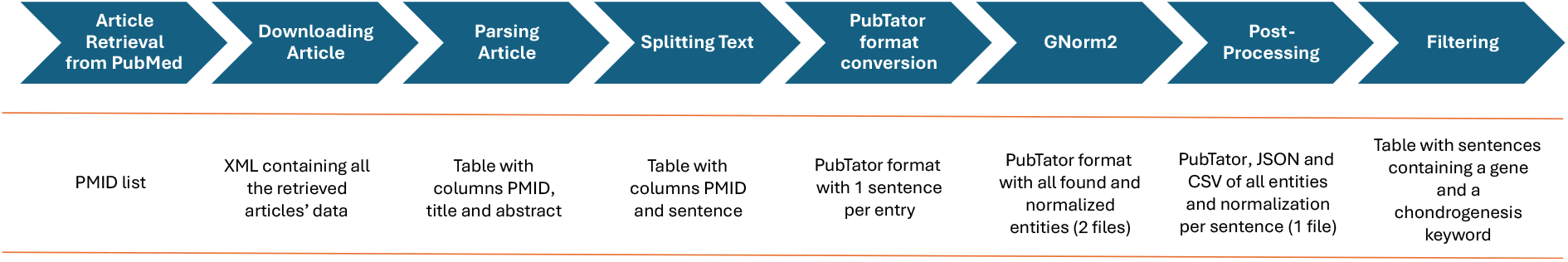
Literature mining and gene annotation workflow. Schematic overview of the text-mining pipeline used to identify and annotate gene mentions from the biomedical literature. Relevant articles were first retrieved from PubMed and downloaded for processing.

This pipeline generated all the datasets used for this study. The specific queries used for extraction are detailed in the dataset section below; the remaining pipeline steps were consistent across all datasets. Entity, abstract, sentence and gene counts at each step are given in Supplementary Table S2.

### Annotation

Two independent curators with backgrounds in molecular biology annotated the gene mentions as *Driver* or *Non-Driver* of chondrogenesis. *Drivers* are genes that the sentence presents as participating in the differentiation of stem cells into chondrocytes. Labels reflect the claim made in the sentence rather than the annotator’s prior knowledge of the gene, so the same gene may receive different labels across sentences. *Non-Drivers* are all remaining gene mentions. The full criteria used to classify the genes into these two categories are detailed in Supplementary Note S1. Discrepancies between curators and genes marked as unclear were resolved by consensus. Inter-annotator agreement over the full annotated set (training, validation and held out) prior to consensus resolution was moderate (Cohen’s κ = 0.57).

### Datasets for Model Development and Evaluation

The training, validation, and LLM prompt development datasets were drawn from a common unlabelled pool built using the pipeline above, restricted to publications up to a cutoff date of 31st of January 2025. The initial training set consisted of 148 sentences selected via a k-medoid greedy algorithm, using SBioBERT (Deka, Jurek-Loughrey, and Deepak 2022) with sentence embeddings as the feature space and cosine similarity as the distance metric. In subsequent iterations, 100 sentences were selected from the unlabelled pool based on the criteria outlined in the Active Learning Loop section. The validation dataset was sampled from the same pool with no PMID-level overlap, and was used to monitor performance, guide hyperparameter decisions, and define the stopping criterion during the active learning loop.

The heldout dataset used the same query and pipeline but was restricted to publications from the 1st of June to the 31st of December 2025 to avoid overlap with the training data. The heldout set was used to compare the fine-tuned PubMedBERT with the other models. It was never used for training, tuning, or model selection for any reported model.

The LLM Prompt Development Dataset was drawn from the final training iteration, enriched with Driver genes to address class imbalance, and used exclusively to refine system and user prompts for LLM-based classification.

All gene instances across these datasets were manually annotated by two curators as *Driver* or *Non-Driver*. Summary statistics for all of them are given in Table 1.

**Table 1.**
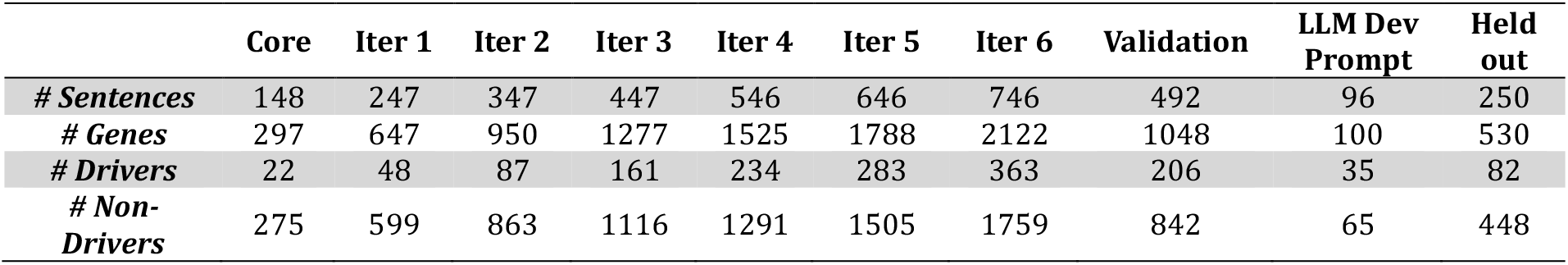
Characteristics of the training and validation dataset used in the active learning framework.

|  | Core | Iter 1 | Iter 2 | Iter 3 | Iter 4 | Iter 5 | Iter 6 | Validation | LLM Dev Prompt | Held out |
| --- | --- | --- | --- | --- | --- | --- | --- | --- | --- | --- |
| # Sentences | 148 | 247 | 347 | 447 | 546 | 646 | 746 | 492 | 96 | 250 |
| # Genes | 297 | 647 | 950 | 1277 | 1525 | 1788 | 2122 | 1048 | 100 | 530 |
| # Drivers | 22 | 48 | 87 | 161 | 234 | 283 | 363 | 206 | 35 | 82 |
| # Non-Drivers | 275 | 599 | 863 | 1116 | 1291 | 1505 | 1759 | 842 | 65 | 448 |

Following model development, the final model was applied to an independent, unannotated application dataset retrieved using the same PubMed search strategy. The dataset comprised publications published between 1st January 2006 and 28th February 2026, with previously labelled gene mentions removed to avoid overlap with the training and evaluation data. The resulting predictions were used to generate a list of candidate regulators.

### Models

#### Input representation

Across all models, gene mentions in the input sentence were masked with the token [GENE], preventing models from memorising gene identities and encouraging them to learn contextual patterns. The gene mention to be classified was delimited by [TARGET] and [/TARGET] to distinguish it from other gene mentions in the same sentence. Each gene mention was treated as an independent instance, so classification was performed at the mention level rather than the gene level.

#### PubMedBERT

Classification was performed using PubMedBERT (Gu *et al*. 2021), specifically the version available on the HuggingFace platform as *microsoft/BiomedNLP-BiomedBERT-base-uncased-abstract-fulltext*. The model was fine-tuned within an active learning framework as a binary classifier to determine whether a gene mention was associated with a driver role in chondrogenesis in the presented text.

#### Baseline Model

To establish a baseline for the gene classification task, we used a logistic regression classifier trained on embeddings from the core training dataset. These embeddings are the [CLS] token of the last hidden layer of the PubMedBERT model, which is a summarised representation of the entire input sequence.

#### Generative Models

We evaluated four models from the Qwen3 (Yang *et al*. 2025) family on the same classification task: a 4B instruction-oriented model (Qwen3-4B-Instruct-2507), a 4B reasoning-oriented model (Qwen3-4B-Thinking-2507), a quantised 30B instruction-oriented model (Qwen3-30B-A3B-Instruct-2507-FP8), and a quantised 30B reasoning-oriented model (Qwen3-30B-A3B-Thinking-2507-FP8); as well as one model from the Llama family, Meta-Llama-3.1-70B-Instruct (AWQ INT4) (Grattafiori *et al*. 2024).

Models were prompted zero- and few-shot with a shared system prompt requiring the models to generate a single-word output, “Yes” or “No”, corresponding to the *Driver* or *Non-Driver* classes, respectively. The full system, zero- and few-shot prompts can be found in the Supplementary Note S2-4. Generation was constrained to a single token, and the class was assigned directly from the generated token. The probability associated with that token was taken as the model’s confidence in the predicted class and used to derive the continuous scores required for AUC-ROC.

Performance of the generative models was evaluated on the held-out dataset, enabling comparison with PubMedBERT. For all models in this comparison, 95% confidence intervals for all reported metrics were estimated via non-parametric bootstrap resampling (10,000 iterations with replacement) of predictions from the heldout dataset.

#### Compute Environment

All models were trained and evaluated on a single NVIDIA L40S GPU (48 GB VRAM) node in an institutional HPC cluster. Because of incompatible dependency requirements across model families, PubMedBERT, the Qwen3 models, and Llama-3.1-70B were each run in separate Python virtual environments, differing principally in their *transformers, torch* and *huggingface* versions. Package versions for each environment are listed in Supplementary Table S4. Hardware was identical across all runs, so differences in compute resources do not confound throughput comparisons.

### Active Learning Loop

The active learning framework was implemented as an iterative process in which the model was progressively improved while the training dataset was incrementally expanded through successive annotation cycles (Figure 2). The loop was terminated once the model reached an AUC-ROC of 0.9 on the validation set, which was set as the stopping criteria.

**Figure 2.**
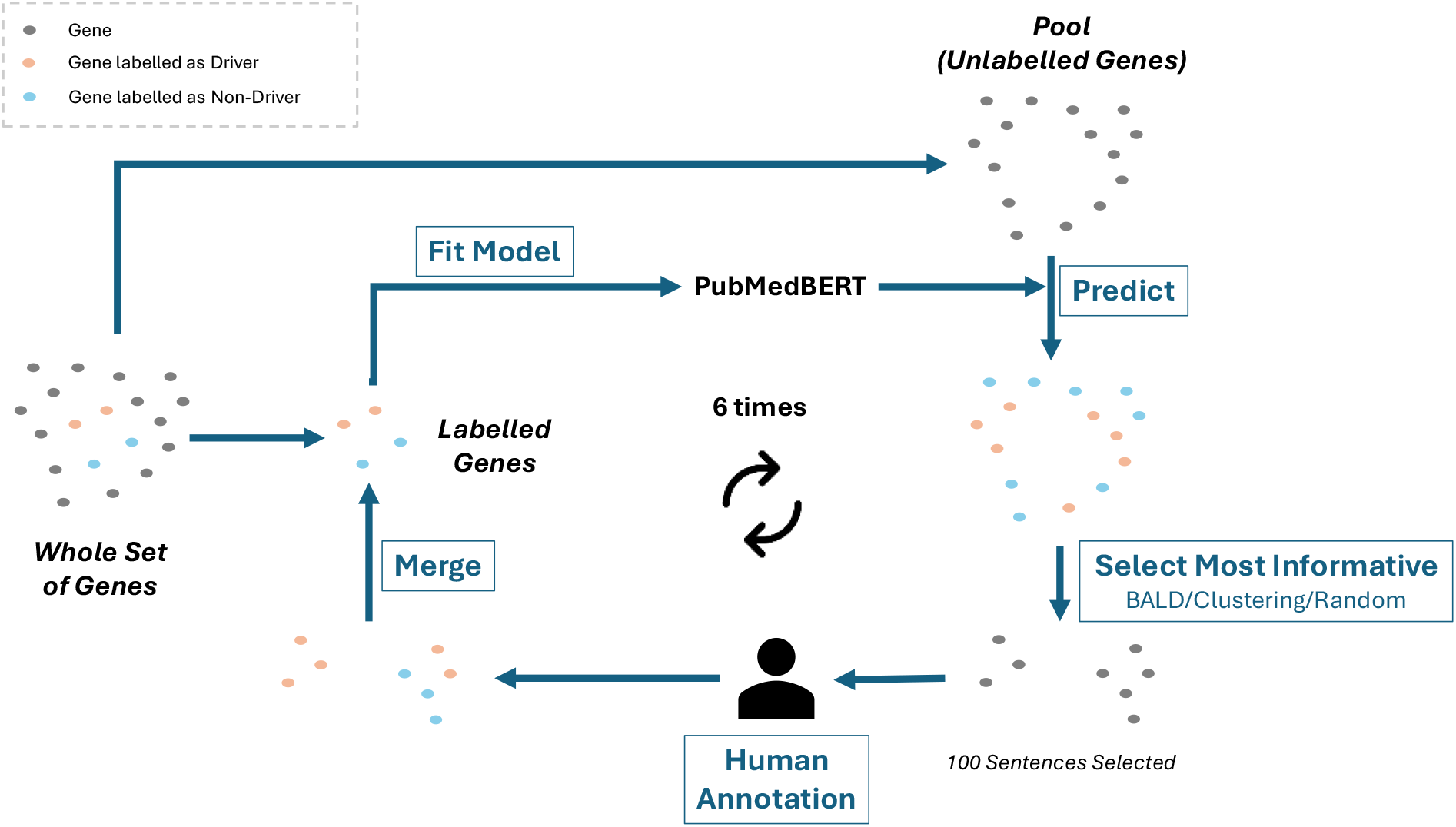
Schematic overview of the active learning loop used in this work. Starting from an initial set of labelled genes, PubMedBERT was fitted and used to predict on the pool of unlabelled genes. The most informative genes were then selected using a combination of BALD, diversity-based clustering, and random sampling, yielding 100 sentences per iteration for human annotation. Newly annotated genes were merged into the labelled set, which was used to refit the model in the next cycle. This loop was repeated six times.

For each iteration, samples were selected using a combination of three strategies: diversity-based sampling through k-medoids clustering, where the medoid and outliers of each cluster were selected; uncertainty-based sampling using Bayesian Active Learning by Disagreement (BALD) (Cao and Tsang 2021) implemented through Monte Carlo dropout over 200 stochastic forward passes (Gal and Ghahramani 2016); and random sampling.

The number of samples chosen by each technique varied across different iterations (Supplementary Table S3), with diversity sampling dominant early and BALD-based uncertainty sampling weighted more heavily as the dataset grew to prioritise more informative samples.

Since the classification model operates at the gene-mention level but annotation was performed on whole sentences, any sentence containing a selected gene was taken as the full sample to annotate. If a selected gene’s sentence had already been selected, it was replaced by the nearest unselected gene mention in embedding space (cosine similarity). This replacement step was repeated until 100 distinct sentences had been collected per iteration.

From iterations 3 to 5, class-balancing strategies were incorporated into the sampling process to address the imbalance of the minority Driver class. These strategies included a weighted variant of BALD, in which samples were up-weighted based on the rarity of their predicted class. This approach follows the class-balancing framework for efficient active learning on imbalanced datasets, as described in (Fairstein *et al*. 2024). Additionally, during model training for these iterations, we used a weighted loss function to further penalise misclassifications of the minority class.

#### Random sampling comparison

To contextualise the gains from active learning, we implemented a random-sampling comparison using the base PubMedBERT model. We randomly selected genes and their associated sentences using the same sentence-selection procedure as in the active learning process. However, for each model training run, we selected the genes independently, without accumulating selections across iterations as in the active learning loop. We selected the same number of sentences as used across the active learning iterations, trained the model on this randomly sampled dataset, and evaluated it on the validation set. To account for variance introduced by the random selection, this procedure was repeated with six random seeds, and the results were averaged across runs. This comparison was conservative: sampling was restricted to the training and held-out sets, since these were the only labelled samples available, rather than to the full unlabelled pool used by the active learning loop. This restricted pool was considerably smaller and less noisy, making random selection an easier task than it would be in a real-world scenario.

#### Metrics

Model performance was evaluated using AUC-ROC, F1-score, precision and recall, with Driver as the positive class. All metrics were computed at the mention level, consistent with the input representation described above. Precision, recall and F1 were calculated at a decision threshold of 0.5.

These metrics were selected to provide a comprehensive view: AUC-ROC for threshold-independent discrimination, F1-score for balancing precision and recall, and precision and recall assessing false-positive and false-negative rates, respectively, which are especially important given the class imbalance in our datasets.

### Downstream Analysis

The production PubMedBERT model was applied to all gene mentions in the application dataset. Mentions with an NCBI Gene identifier provided by GNorm2 during the data extraction pipeline were retained, and those left unnormalised were, where possible, resolved using the Python package *mygene* (v3.2.2). Mentions that remained unresolved were excluded.

Normalised mentions were filtered to human genes (taxid 9606) and aggregated by a disjunctive rule. If any mention of a gene was classified as Driver, that gene was retained as a candidate regulator.

Candidates were compared with human genes annotated with *chondrocyte differentiation* (GO:0002062). Annotations were taken from org.Hs.eg.db (v3.22.0) and the ontology from GO.db (v3.22.0) (Ashburner *et al*. 2000, The Gene Ontology Consortium 2026), with terms expanded via GOALL to include descendant annotations. The comparison used genes classified as *Driver* by the fine-tuned PubMedBERT model and subsequently normalised, as described in the previous section. The same procedure was applied to the parent term cartilage development (GO:0051216), excluding genes annotated under GO:0002062 or its descendants. Analyses were performed in R 4.5.2.

## Results

### A small annotation budget is sufficient to achieve a classifier that ranks candidate regulators reliably

Extracting regulators of chondrogenesis from the literature requires a classification model, but manual sentence annotation is time-consuming. To reduce this cost, we used an active learning framework to select the training set, which ran for six iterations (Figure 2). An initial core set of 148 sentences was selected from the unlabelled pool using a k-medoids greedy algorithm and annotated by two curators, yielding 297 gene instances. At each subsequent iteration, 100 sentences were selected from the unlabelled pool based on uncertainty, diversity, and random sampling. Their gene mentions were annotated and added to the training set. AUC-ROC, a threshold-independent measure of class separability, guided all decisions during the learning framework, and we used an AUC-ROC exceeding 0.9 as the stopping criterion, so that the loop terminated on a performance threshold rather than an arbitrary iteration count.

The composition of the training dataset and model performance were evaluated across iterations (Figure 3). The model trained on the core set achieved an AUC-ROC of 0.68 but assigned every validation gene to the Non-Driver class, resulting in a precision, recall, and F1-score of 0. Iterations 1 and 2, using the unweighted hybrid strategy, improved AUC-ROC slowly, to 0.74 and 0.78, while recall and F1 remained low. Sampling selected few positive examples at each step (26 and 39 Driver genes), leaving the class imbalance largely unchanged and, we hypothesised, limiting the rate of improvement.

**Figure 3.**
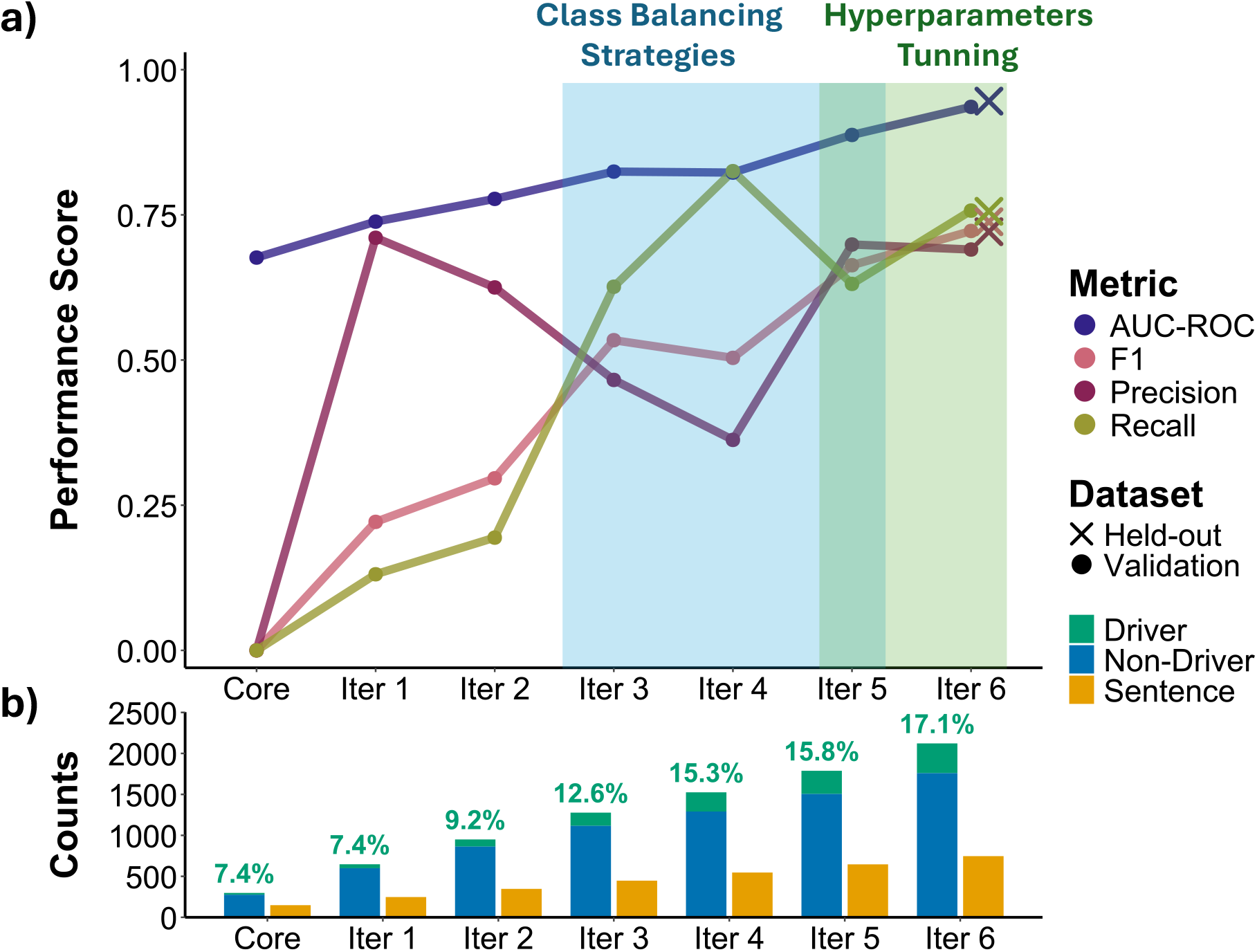
Performance of the model across active learning iterations and training data composition. a) AUC-ROC, F1-score, precision, and recall performance scores are shown for all the iterations of the active learning loop. Shaded boxes indicate iterations where additional modelling or sampling strategies were applied. b) The number of gene and sentence instances in the training dataset at each iteration. The gene bars are split by the amount of each class the training data has, and on top of the bar, the percentage of gene mentions classified as Driver.

To accelerate improvement, we introduced two class-balancing strategies at iteration 3. The first was a class-weighted loss applied during training to improve the model’s discrimination between the minority class (Driver) and other classes. The second was class weighting applied at the sampling stage, which adjusted the existing uncertainty sampling criteria to favour genes predicted as Driver. Applying these strategies, recall rose from 0.30 to 0.53, F1 from 0.26 to 0.53, and AUC-ROC reached 0.82, although precision declined, reflecting more false positives (Figure 3a).

Performance then plateaued at iteration 4 (AUC-ROC 0.82, F1 0.50), indicating that further training data alone was no longer improving performance. From this point, we introduced a grid search over epochs, learning rate, batch size, warmup steps, weight decay, and use of class-weighted loss (Supplementary Table S5). At iteration 5, we observed the largest increase in AUC-ROC to 0.89. Other metrics also increased: F1 to 0.66 and precision to 0.70, reversing the downward trend seen from iterations 2 to 4, while recall fell to 0.63.

In iteration 6, an initial training run with class-weighted loss resulted in a sharp drop in all metrics (AUC-ROC 0.70, F1-score 0.44, precision 0.36, recall 0.59), suggesting the majority class was being over-penalised. Removing class-weighted loss recovered performance (AUC-ROC 0.89, F1 0.60), and hyperparameter tuning under this configuration reached an AUC-ROC of 0.94, meeting the stopping criterion.

As the validation dataset was used to guide model selection, balancing strategies and hyperparameter tuning throughout the loop, it was no longer an independent estimate of performance. We therefore evaluated the iteration 6 model on the held-out dataset, obtaining an AUC-ROC of 0.95, an F1-score of 0.74, a precision of 0.72, and a recall of 0.76 (Figure 3, crosses). The performance was consistent with the validation results, indicating that the gains accumulated over the loop generalise to unseen literature despite the validation set’s role in development. In total, 746 sentences were used for the last trained model in iteration 6, corresponding to 2,122 gene instances, of which 363 were labelled Driver and 1,759 Non-Driver (Figure 3b).

To test whether the performance gains came from the selection strategy or from the added data alone, we compared the trajectory against random draws from the labelled pool, matched for sentence count (Figure 4). AUC-ROC and F1 were similar under both. The comparison was limited, however, because the pool was assembled by the active learning run itself, so random draws from it are already enriched for informative sentences. Replicating the active learning trajectory across seeds was not attempted, as each seed’s selection of sentences would require separate annotation. Class composition (Supplementary Figure S1) differed between the two approaches as Drivers rose from 7.4% to 17.1% under active learning and from 11.5% to 15.8% under random selection (mean of 6 seeds), the lower starting point reflecting a core set selected for diversity alone, without regard to class balance.

**Figure 4.**
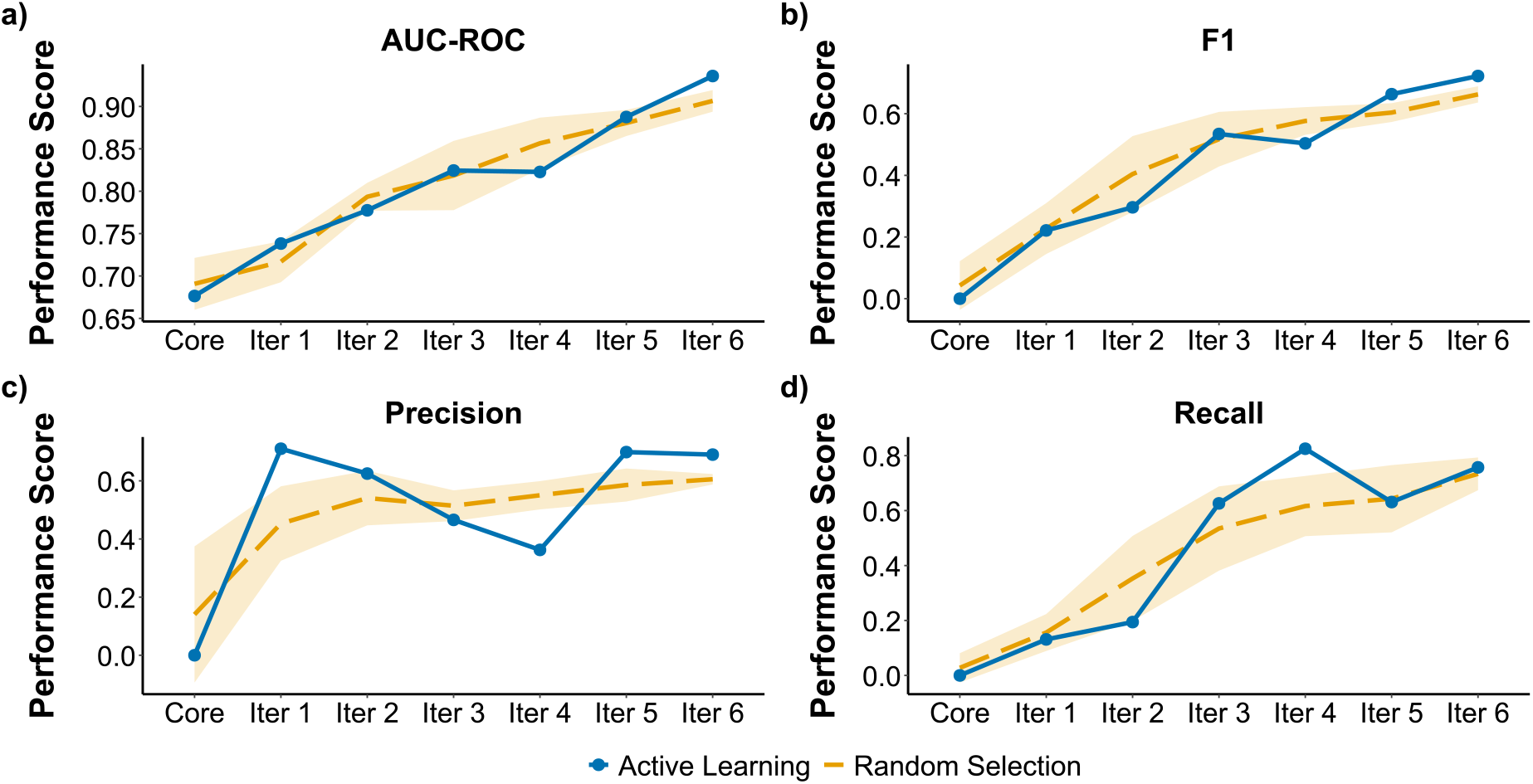
Comparison of active learning and random sampling strategies for gene regulator classification across annotation iterations. (a–d) AUC-ROC, F1, Precision, and Recall were evaluated on the validation dataset from the initial seed set (“Core”) across six annotation iterations. The active learning strategy is represented as a solid blue line, whereas the random sampling baseline is represented as a dashed orange line. Shaded regions indicate the 95% confidence interval across six random seeds.

### Fine-tuned PubMedBERT matches the gLLMs in performance at a fraction of the computational cost

After the active learning loop was completed, PubMedBERT was retrained on the combined training and validation datasets to produce the final production model. The held-out dataset was not used for training, tuning or model selection at any stage. We compared PubMedBERT against a logistic regression baseline and open-weight generative large language models (gLLMs): Qwen3 at 4B and 30B parameters, each in instruction-tuned and reasoning variants, and Llama 3.1 70B instruction-tuned. All models ran under the same hardware conditions.

Performance scores (AUC-ROC, F1, precision, and recall) were calculated for each model, along with the computational time for model training and prediction on the held-out dataset, with 95% confidence intervals (Table 2). Only few-shot prompting (11 examples) is reported for the gLLMs, as few-shot prompting performed comparably to zero-shot prompting across all models. Results for both prompting strategies are given in Supplementary Table S6.

**Table 2.** Model performance, configuration, and computational requirements for gene mention classification.

| Model | Parameters | Tuning Variant | Training Time (s) | Average Gene Inference Time (ms) | Projected corpus runtime (67111 genes) | Precision | Recall | F1 | AUC-ROC |
| --- | --- | --- | --- | --- | --- | --- | --- | --- | --- |
| **Logistic Regression** | - | - | 36.86 | 0.003 | 0.2s | 0.35<br>[0.09, 0.64] | 0.09<br>[0.02, 0.17] | 0.14<br>[0.03, 0.26] | 0.69<br>[0.61, 0.76] |
| **PubMedBERT** | 109M | - | 88.58 | 4.1 | 4.6min | 0.69<br>[0.56, 0.82] | 0.66<br>[0.53, 0.78] | 0.67<br>[0.56, 0.77] | 0.93<br>[0.89, 0.96] |
| **Qwen3** | 4B | Instruction | - | 890 | 16.59h | 0.43<br>[0.34, 0.52] | 0.68<br>[0.57, 0.8] | 0.52<br>[0.43, 0.61] | 0.86<br>[0.8, 0.9] |
| **Qwen3** | 30B | Instruction | - | 1,098 | 20.47h | 0.42<br>[0.33, 0.52] | 0.85<br>[0.76, 0.94] | 0.57<br>[0.47, 0.65] | 0.91<br>[0.88, 0.94] |
| **Llama3.1** | 70B | Instruction | - | 1,822 | 1.42d | 0.43<br>[0.35, 0.52] | 0.93<br>[0.85, 0.99] | 0.59<br>[0.5, 0.68] | 0.94<br>[0.91, 0.96] |
| **Qwen3** | 4B | Reasoning | - | 118,746 | 92.24d | 0.64<br>[0.52, 0.76] | 0.74<br>[0.63, 0.85] | 0.69<br>[0.59, 0.78] | 0.85<br>[0.79, 0.91] |
| **Qwen3** | 30B | Reasoning | - | 137,321 | 106.66d | 0.68<br>[0.57, 0.78] | 0.78<br>[0.67, 0.88] | 0.73<br>[0.63, 0.81] | 0.94<br>[0.91, 0.97] |

All language models outperformed the logistic regression baseline. Among the Qwen3 models, the reasoning-oriented variants outperformed the instruction-tuned ones in precision (0.64 and 0.68 at 4B and 30B, versus 0.43 and 0.42) and F1 (0.69 and 0.73, versus 0.52 and 0.57), consistent with the task requiring interpretation of experimental context and the association of evidence with specific gene mentions. Parameter size in this family of models has a more limited effect: recall and AUC-ROC improved with scale in both variants, whereas precision improved only in the reasoning ones.

Llama 3.1 70B extended this pattern beyond the Qwen3 family. Across the three instruction-tuned models, precision remained consistent regardless of scale (0.43, 0.42 and 0.43), and F1 stayed below both reasoning-oriented variants, while recall and AUC-ROC increased with scale. Llama 3.1 70B reached the highest recall of any model evaluated (0.93) and an AUC-ROC of 0.94, matching Qwen3-30B reasoning.

On the same held-out dataset, the fine-tuned PubMedBERT model achieved an AUC-ROC of 0.93, marginally below the top two gLLMs, and the highest precision of any model evaluated (0.69), at a recall of 0.66. Although PubMedBERT and Llama 3.1 70B ranked genes with comparable accuracy, they classified differently at their default thresholds. Llama 3.1 70B recovered almost all Driver genes (recall 0.93) at low precision (0.43), flagging many false positives, whereas PubMedBERT was more balanced and achieved the higher F1 of the two (0.67 versus 0.59).

Differences in computational cost, specifically inference time, were considerably larger than the differences in performance. Per-mention gene inference time was 133× higher for Qwen3-4B reasoning than its instruction-tuned counterpart, and 125× higher at 30B. Projected across the 67,111 genes mentions in the full PubMed corpus, the reasoning models would require more than three months of continuous inference, compared to ∼1.5 days for Llama 3.1 70B, under a day for the Qwen3 instruction-tuned models, and just over 4.5 minutes for PubMedBERT. The training cost of the last model, 88.6 seconds, is negligible.

These results demonstrate that a domain-specific encoder fine-tuned on 1,238 annotated sentences performs comparably to the strongest open-weight models evaluated here, while remaining the only approach among them that can feasibly be applied to the full corpus under a single-GPU constraint.

### Candidate regulators recover most of the annotated cartilage development gene set and substantially extend it

Applying the production PubMedBERT classifier to the application dataset (67,111 gene mentions) took 5 minutes and 34 seconds, of which 1 minute and 57 seconds was spent on prediction, a prediction time lower than the calculated projected time reported for this corpus (Table 2). The model classified 10,261 Driver and 56,850 Non-Driver gene mentions.

We retained human gene mentions normalised to Entrez identifiers by GNorm2 or the R package mygene, discarding those that neither method could normalise. Aggregating to gene level, where a gene is treated as a candidate chondrogenesis driver if at least one of its mentions is classified as such, gave 2,776 unique genes, of which 1,128 (40.6%) were classified as candidate drivers. Supporting mentions per candidate ranged from 1 to 829 (median 2, mean 6.56), with 427 (37.9%) supported by a single gene mention (Supplementary Figure 2).

To evaluate how well these predictions recover curated knowledge, we compared the 1,128 candidates against the 119 human genes annotated in the GO term chondrocyte differentiation (GO:0002062) and its descendants. The model recovered 94 of these annotated genes (79%) (Figure 5). Furthermore, an over-representation analysis of the full regulator candidate set using GO Biological Process terms showed to align with the target domain, with terms such as cartilage and connective tissue development among the top 10 significantly enriched terms (Supplementary Figure S3).

**Figure 5.**
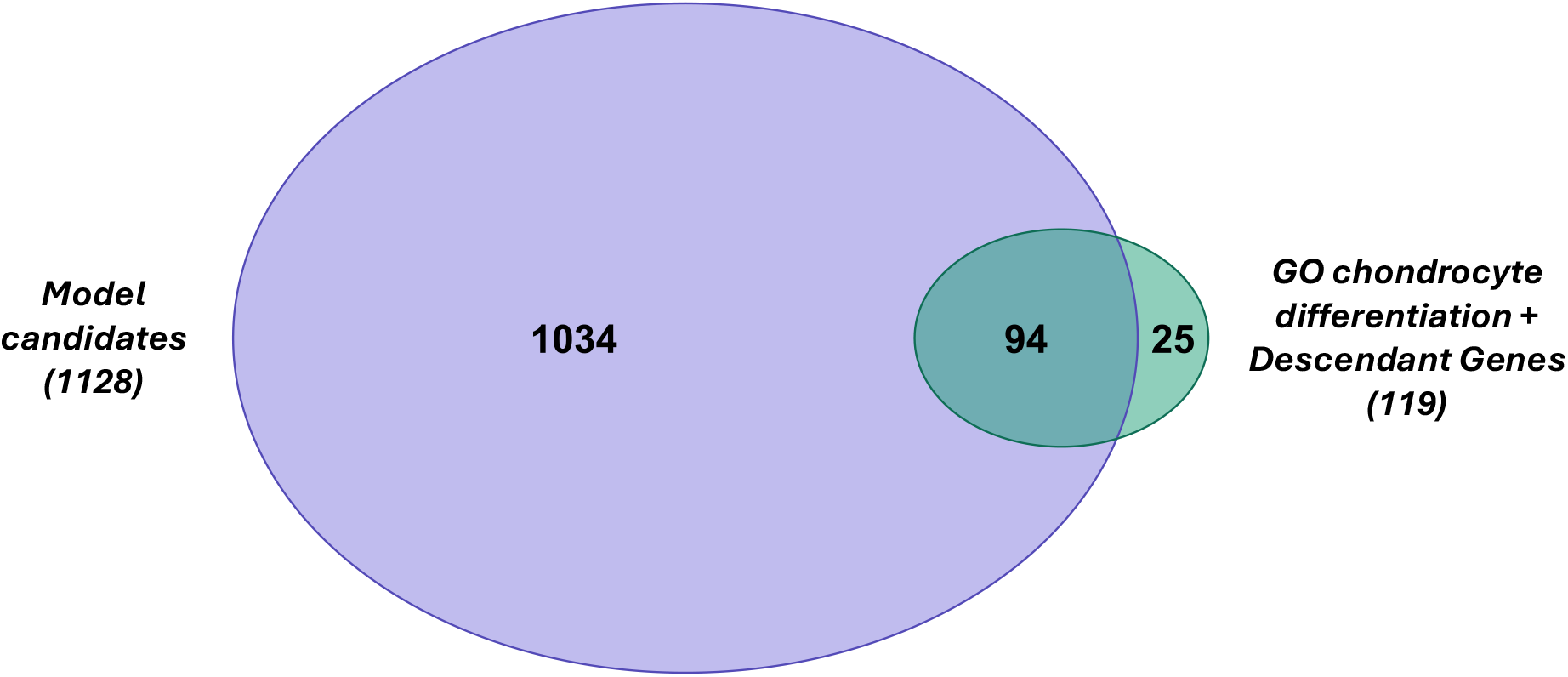
Overlap between predicted regulator candidates and GO cartilage development annotations. Venn diagram comparing the 1,128 predicted chondrogenesis regulators against 119 genes annotated under GO:0002062 (chondrocyte differentiation) and its descendants.

Of the 25 not recovered, 14 were present in the PubMed corpus but were not classified as drivers. Not all of these represented misclassification of the gene mention, since a gene can be discussed in a chondrogenic context without a regulatory relationship being asserted, for example, as a biomarker, which our guidelines treat as *Non-Driver*. The remaining 11 genes were absent from the normalised gene set, potentially reflecting their absence from the corpus, species filtering, or normalisation failure.

The remaining 1034 candidate regulators were not annotated to GO:0051216 or its descendants. The set includes genes with well-documented roles in cartilage development like Transforming Growth Factor Beta 3 (TGFB3), Insulin-like Growth Factor 1 (IGF1), and Tumour Necrosis Factor (TNF) (Schmidt, Chen, and Lynch 2006, Wehling *et al*. 2009, Du *et al*. 2023). The oldest evidence in the supporting studies, dating to 2006, suggests the gap is unlikely to be solely a matter of annotation lag. Of these 1034 genes that were not covered by chondrocyte differentiation, 53 fall under the related term cartilage development (GO:0051216), parent of GO:0002062, or its other descendants.

## Discussion

Annotated datasets are rarely available for specialised text extraction tasks, and manually curating labelled data is costly. We address this scarcity in the context of identifying drivers of chondrogenesis, comparing a domain-specific encoder trained against open-weight LLMs. We developed a fine-tuned PubMedBERT classification model that achieved an AUC-ROC of 0.93, using 1,238 labelled sentences, and required 88.6 seconds of training and 1 minute 57 seconds to predict the full 67,111-gene corpus.

We trained this classification model with 3,170 genes across 1,238 labelled sentences. Previous studies in Bio-NLP active learning have reported budgets ranging from about 500 to over 20,000 labelled samples (Zhou, He, and Kwoh 2006, Chen Y, Mani, and Xu 2012, Guo *et al*. 2013, Giles *et al*. 2020, Can izares-Dí az *et al*. 2021, Nachtegael *et al*. 2024). Our dataset of 1,238 samples falls toward the lower end of this range. There are two reasons why our gene count is relatively high despite the smaller sample size. First, driver genes are rare in the data, leading to many non-driver labels even with minimised labelling strategies. Second, since annotation is done at the sentence level and sentences often mention multiple genes, the number of labelled genes increases faster than the number of sentences. More aggressive class balancing or earlier uncertainty sampling might have reduced this count.

AUC-ROC and F1-score were similar under active learning random selection, so we cannot attribute the observed gains to the sampling strategy. The random baseline was drawn from the labelled dataset assembled by the active learning run itself, together with the held-out set, rather than from an independent sample of the full corpus, so it is already enriched with informative sentences. Nevertheless, the total annotation effort remained well below what is typically required for fully supervised biomedical NLP work (Campos *et al*. 2012, Campos, Matos, and Oliveira 2015, Li *et al*. 2021, Hu *et al*. 2022, Nachtegael *et al*. 2024), and was sufficient to produce a classifier that generalised to unseen literature.

We additionally investigated whether the annotation effort was justified, given that generative models require no task-specific labelling. Since the answer depends on which models a group can realistically run, we restricted the comparison to open-weight models that fit on a single GPU.

Our fine-tuned PubMedBERT performs comparably to open-weight generative models that are one to two orders of magnitude larger. Which metric matters most depends on how the classifier is used. If the models are used directly, the positive class must be trustworthy, as every false positive costs the curator time to reject. On precision at default thresholds, PubMedBERT was among the strongest models evaluated (0.69). If the model serves as a prioritisation step, producing a ranked list curated from the top down and with no threshold applied, separability is the relevant measure. Here, PubMedBERT’s AUC-ROC of 0.93 is on par with 0.94 for both Qwen3-30B-Reasoning and Llama 3.1 70B. With performance comparable between models, cost becomes the deciding factor. Our classification target is the full retrieved corpus of 67,111 gene mentions, and this may not be a one-off cost, as any revision to the labelling criteria would require reclassifying the entire corpus. PubMedBERT’s projected 4.6 minutes per pass makes such reclassification trivial, whereas 1.42 days for Llama 3.1 70B and 106.66 days for Qwen3-30B-Reasoning place a practical ceiling on how often it can be done. However, these times apply to our single-GPU setup and may vary with different hardware or configurations.

Of the two generative models matching PubMedBERT on AUC-ROC, only Llama 3.1 70B is cheap enough for repeated reclassification. Nevertheless, its precision of 0.43 places it with the instruction-tuned variants rather than the reasoning models, despite being more than twice the size of the largest Qwen3 tested. When we group the models by variant, the reasoning variants outperform their instruction-tuned counterparts of the same size by roughly 0.25 in precision, but at 125–133 times the inference cost. However, with only two model families, this pattern cannot be assumed to be general, though it suggests a larger instruction-tuned model is not a path to reasoning-level precision. These findings were determined by our choice to compare only open-weight models that can run on a single GPU. Frontier commercial models may outperform every model evaluated here, but access is typically constrained by cost, rate limits, and, for groups working with clinical or unpublished text, by restrictions on sending it to external APIs.

Applying the production PubMedBERT model to the full corpus returned a candidate set of regulators of chondrogenesis, 86% of which lack a cartilage development annotation in GO. This is not a straightforward false-positive rate, since GO coverage is incomplete: an absent annotation may reflect no true role, a classifier error, or a genuine regulatory role not yet curated. This candidate set presents two limitations. First, 427 of the 1,128 candidates (37.9%) rest on a single mention classified as a Driver, offering no redundancy against gene misclassification or normalisation errors. The ranked list should therefore be read as a prioritisation for curation, with single-mention candidates checked against their source sentences before any experimental follow-up. Second, the entity normalisation process removed a considerable number of gene mentions and optimising that process could yield more candidate regulators.

Examining other generative models such as Mistral (Jiang *et al*. 2023) and Gemma (Team *et al*. 2024) would establish whether the instruction/reasoning pattern generalises across families. During prompt development, reasoning models sometimes correctly identified driver genes based on phenotypic evidence not in our guidelines, suggesting a potential role in refining annotation criteria or flagging curator disagreement, despite limited applicability at scale. We compared two options, building a labelled dataset for fine-tuning or using gLLMs directly, but a third exists: pseudo-labelling with a gLLM and fine-tuning the encoder on those labels, thereby removing manual annotation entirely (Pangakis and Wolken 2024, Zhao Y and Goto 2025). Qwen3-30B-Reasoning reached an F1 of 0.73 here; this is a plausible future direction. Another possible improvement to the study would be enhancing the random baseline. Currently, it samples sentences from the already-annotated candidate-mention pool used for active learning, rather than from the entire unlabelled corpus. Creating a baseline from the full corpus would provide a more rigorous evaluation of the annotation strategy’s effectiveness.

## Conclusion

We fine-tuned PubMedBERT to identify regulators of chondrogenesis in the literature, yielding a robust classification model trained on 1,238 labelled sentences selected via active learning. Under a single-GPU, local-inference constraint, this model matched open-weight generative models that are one to two orders of magnitude larger in both precision and AUC-ROC and classifies the full corpus in less than 6 minutes, compared with projected runtimes of days to months for generative models.

Applied at the corpus scale, the model returned 1,128 candidate regulators, 92% of which were not annotated for chondrocyte differentiation in GO. This ranked list is intended to seed manual curation of a high-confidence regulator set, which could be used as labels for different machine learning methods, such as graph-based models over biological networks enriched with chondrogenesis-specific omics data (Soul and Young 2026).

As the pipeline is not specific to chondrogenesis, it should transfer to other tasks where the positive class is rare, and a domain-specific encoder can be used, though whether a comparable annotation budget suffices in these settings remains to be tested.

## Acknowledgements

The Authors acknowledge use of the HPC cluster provided by the Digital Research Infrastructure Team (TIED), University of Liverpool.

During the preparation of this manuscript, the authors used Claude (Anthropic) to assist with language editing and, on occasion, with debugging and improving existing research code. All AI-assisted suggestions were reviewed, tested, and verified by the authors. The authors remain fully responsible for the final code, analyses, results, and manuscript.

## References

Agrawal M, Hegselmann S, Lang H et al. Large Language Models are Few-Shot Clinical Information Extractors, arXiv:2205.12689. Preprint, arXiv, 30 Nov. 2022. 10.48550/arXiv.2205.12689.

Ash JT, Zhang C, Krishnamurthy A et al. Deep Batch Active Learning by Diverse, Uncertain Gradient Lower Bounds, paper delivered at International Conference on Learning Representations. 25 Sept. 2019. https://openreview.net/forum?id=ryghZJBKPS (30 May 2025, date last accessed).

Ashburner M, Ball CA, Blake JA et al. Gene Ontology: tool for the unification of biology. Nat Genet 2000;25(1):25–9. 10.1038/75556.

Azadifar S, Ahmadi A. A novel candidate disease gene prioritization method using deep graph convolutional networks and semi-supervised learning. BMC Bioinformatics 2022;23(1):422. 10.1186/s12859-022-04954-x.

Bada M, Eckert M, Evans D et al. Concept annotation in the CRAFT corpus. BMC Bioinformatics 2012;13(1):161. 10.1186/1471-2105-13-161.

Balachandran S, Prada-Medina CA, Mensah MA et al. STIGMA: Single-cell tissue-specific gene prioritization using machine learning. Am J Hum Genet 2024;111(2):338–49. 10.1016/j.ajhg.2023.12.011.

Beltagy I, Lo K, Cohan A. SciBERT: A Pretrained Language Model for Scientific Text, arXiv:1903.10676. Preprint, arXiv, 10 Sept. 2019. 10.48550/arXiv.1903.10676.

Ben Abacha A, Zweigenbaum P. Automatic extraction of semantic relations between medical entities: a rule based approach. J Biomed Semant 2011;2(5):S4. 10.1186/2041-1480-2-S5-S4.

Blanchard AE, Gao S, Yoon HJ et al. A Keyword-Enhanced Approach to Handle Class Imbalance in Clinical Text Classification. IEEE J Biomed Health Inform 2022;26(6):2796–803. 10.1109/JBHI.2022.3141976.

Boeuf S, Richter W. Chondrogenesis of mesenchymal stem cells: role of tissue source and inducing factors. Stem Cell Res Ther 2010;1(4):31. 10.1186/scrt31.

Bose P, Srinivasan S, Sleeman WC et al. A Survey on Recent Named Entity Recognition and Relationship Extraction Techniques on Clinical Texts. Appl Sci 2021;11(18):8319. 10.3390/app11188319.

Bumgardner VKC, Mullen A, Armstrong SE et al. Local Large Language Models for Complex Structured Tasks. AMIA Summits Transl Sci Proc 2024;2024:105–14.

Campos D, Matos S, Lewin I et al. Harmonization of gene/protein annotations: towards a gold standard MEDLINE. Bioinformatics 2012;28(9):1253–61. 10.1093/bioinformatics/bts125.

Campos D, Matos S, Oliveira JL. A document processing pipeline for annotating chemical entities in scientific documents. J Cheminformatics 2015;7(1):S7. 10.1186/1758-2946-7-S1-S7.

Canizares-Díaz H, Piad-Morffis A, Estevez-Velarde S et al. Active Learning for Assisted Corpus Construction: A Case Study in Knowledge Discovery from Biomedical Text. In: Mitkov R, Angelova G (eds), Proceedings of the International Conference on Recent Advances in Natural Language Processing (RANLP 2021). Held Online: INCOMA Ltd., 2021, 216–25. https://aclanthology.org/2021.ranlp-1.26/ (27 Apr. 2026, date last accessed).

Cao X, Tsang IW. Bayesian Active Learning by Disagreements: A Geometric Perspective, arXiv:2105.02543. Preprint, arXiv, 6 May 2021. 10.48550/arXiv.2105.02543.

Chen M, Jiang Z, Zou X et al. Advancements in tissue engineering for articular cartilage regeneration. Heliyon 2024;10(3):e25400. 10.1016/j.heliyon.2024.e25400.

Chen Y, Mani S, Xu H. Applying active learning to assertion classification of concepts in clinical text. J Biomed Inform 2012;45(2):265–72. 10.1016/j.jbi.2011.11.003.

Cheung K, Barter MJ, Falk J et al. Histone ChIP-Seq identifies differential enhancer usage during chondrogenesis as critical for defining cell-type specificity. FASEB J 2020;34(4):5317–31. 10.1096/fj.201902061RR.

Cohn DA, Ghahramani Z, Jordan MI. Active Learning with Statistical Models. J Artif Intell Res 1996;4:129–45. 10.1613/jair.295.

Collins C, Baker S, Brown J et al. Text mining for contexts and relationships in cancer genomics literature. Bioinformatics 2024;40(1):btae021. 10.1093/bioinformatics/btae021.

Collins C, Fytas P, Karadeniz I et al. BioTriplex: a full-text annotated corpus for fine-tuning language models in gene-disease relation extraction tasks. Bioinformatics 2026;42(2):btag037. 10.1093/bioinformatics/btag037.

De Angeli K, Gao S, Danciu I et al. Class imbalance in out-of-distribution datasets: Improving the robustness of the TextCNN for the classification of rare cancer types. J Biomed Inform 2022;125:103957. 10.1016/j.jbi.2021.103957.

Deka P, Jurek-Loughrey A, Deepak. Unsupervised Keyword Combination Query Generation from Online Health Related Content for Evidence-Based Fact Checking. 23rd Int Conf Inf Integr Web Intell (New York, NY, USA) iiWAS2021, 30 Dec. 2022:267–77. 10.1145/3487664.3487701.

Doucet P, Estermann B, Aczel T et al. Bridging Diversity and Uncertainty in Active learning with Self-Supervised Pre-Training, arXiv:2403.03728. Preprint, arXiv, 17 Jan. 2025. 10.48550/arXiv.2403.03728.

Du X, Cai L, Xie J et al. The role of TGF-beta3 in cartilage development and osteoarthritis. Bone Res 2023;11(1):2. 10.1038/s41413-022-00239-4.

Ein-Dor L, Halfon A, Gera A et al. Active Learning for BERT: An Empirical Study. In: Webber B, Cohn T, He Y et al. (eds), Proceedings of the 2020 Conference on Empirical Methods in Natural Language Processing (EMNLP). Online: Association for Computational Linguistics, 2020, 7949–62. 10.18653/v1/2020.emnlp-main.638.

Fairstein Y, Kalinsky O, Karnin Z et al. Class Balancing for Efficient Active Learning in Imbalanced Datasets. In: Henning S, Stede M (eds), Proceedings of the 18th Linguistic Annotation Workshop (LAW-XVIII). St. Julians, Malta: Association for Computational Linguistics, 2024, 77–86. 10.18653/v1/2024.law-1.8.

Fundel K, Ku ffner R, Zimmer R. RelEx—Relation extraction using dependency parse trees. Bioinformatics 2007;23(3):365–71. 10.1093/bioinformatics/btl616.

Gal Y, Ghahramani Z. Dropout as a Bayesian approximation: representing model uncertainty in deep learning. Proc 33rd Int Conf Int Conf Mach Learn -Vol 48 (New York, NY, USA) ICML’16, 19 June 2016:1050–9.

Giles O, Karlsson A, Masiala S et al. Optimising biomedical relationship extraction with BioBERT. Preprint, bioRxiv, 1 Sept. 2020, 2020.09.01.277277. 10.1101/2020.09.01.277277.

Grattafiori A, Dubey A, Jauhri A et al. The Llama 3 Herd of Models, arXiv:2407.21783. Preprint, arXiv, 23 Nov. 2024. 10.48550/arXiv.2407.21783.

Gu Y, Tinn R, Cheng H et al. Domain-Specific Language Model Pretraining for Biomedical Natural Language Processing. ACM Trans Comput Healthc 2021;3(1):2:1-2:23. 10.1145/3458754.

Guo Y, Silins I, Stenius U et al. Active learning-based information structure analysis of full scientific articles and two applications for biomedical literature review. Bioinformatics 2013;29(11):1440–7. 10.1093/bioinformatics/btt163.

He T, Zhang S, Xin J et al. An Active Learning Approach with Uncertainty, Representativeness, and Diversity. Sci World J 2014;2014(1):827586. 10.1155/2014/827586.

Hu Y, He H, Chen Z et al. A Unified Model Using Distantly Supervised Data and Cross-Domain Data in NER. Comput Intell Neurosci 2022;2022(1):1987829. 10.1155/2022/1987829.

Jiang AQ, Sablayrolles A, Mensch A et al. Mistral 7B, arXiv:2310.06825, version 1. Preprint, arXiv, 10 Oct. 2023. 10.48550/arXiv.2310.06825.

Jumaa NF, Razmara J, Parvizpour S et al. Hybrid deep learning models for text-based identification of gene-disease associations. BioImpacts 2025;15(1):31226–31226. 10.34172/bi.31226.

Kanakarajan K raj, Kundumani B, Sankarasubbu M. BioELECTRA:Pretrained Biomedical text Encoder using Discriminators. In: Demner-Fushman D, Cohen KB, Ananiadou S et al. (eds), Proceedings of the 20th Workshop on Biomedical Language Processing. Online: Association for Computational Linguistics, 2021, 143–54. 10.18653/v1/2021.bionlp-1.16.

Kilicoglu H, Rosemblat G, Fiszman M et al. Broad-coverage biomedical relation extraction with SemRep. BMC Bioinformatics 2020;21(1):188. 10.1186/s12859-020-3517-7.

Koyabu S, Phan TTT, Ohkawa T. Extraction of Protein-Protein Interaction from Scientific Articles by Predicting Dominant Keywords. BioMed Res Int 2015;2015(1):928531. 10.1155/2015/928531.

Labrak Y, Rouvier M, Dufour R. A Zero-shot and Few-shot Study of Instruction-Finetuned Large Language Models Applied to Clinical and Biomedical Tasks, arXiv:2307.12114. Preprint, arXiv, 9 June 2024. 10.48550/arXiv.2307.12114.

Le DH. Machine learning-based approaches for disease gene prediction. Brief Funct Genomics 2020;19(5–6):350–63. 10.1093/bfgp/elaa013.

Li D, Xiong Y, Hu B et al. Drug knowledge discovery via multi-task learning and pre-trained models. BMC Med Inform Decis Mak 2021;21(9):251. 10.1186/s12911-021-01614-7.

Luo R, Sun L, Xia Y et al. BioGPT: generative pre-trained transformer for biomedical text generation and mining. Brief Bioinform 2022;23(6):bbac409. 10.1093/bib/bbac409.

Mehryary F, Nastou K, Ohta T et al. STRING-ing together protein complexes: corpus and methods for extracting physical protein interactions from the biomedical literature. Bioinformatics 2024;40(9):btae552. 10.1093/bioinformatics/btae552.

Moreau Y, Tranchevent LC. Computational tools for prioritizing candidate genes: boosting disease gene discovery. Nat Rev Genet 2012;13(8):523–36. 10.1038/nrg3253.

Munro R. Combining uncertainty sampling and diversity sampling. In: Human-in-the-Loop Machine Learning. n.p.: Manning Publications, 2021. https://www.oreilly.com/library/view/human-in-the-loop-machine-learning/9781617296741/OEBPS/Text/05.htm.

Nachtegael C, De Stefani J, Cnudde A et al. DUVEL: an active-learning annotated biomedical corpus for the recognition of oligogenic combinations. Database 2024;2024:baae039. 10.1093/database/baae039.

Neumann M, King D, Beltagy I et al. ScispaCy: Fast and Robust Models for Biomedical Natural Language Processing. Proc 18th BioNLP Workshop Shar Task 2019:319–27. 10.18653/v1/W19-5034.

Nezamuldeen L, Jafri MS. Protein–Protein Interaction Network Extraction Using Text Mining Methods Adds Insight into Autism Spectrum Disorder. Biology 2023;12(10):1344. 10.3390/biology12101344.

Pangakis N, Wolken S. Knowledge Distillation in Automated Annotation: Supervised Text Classification with LLM-Generated Training Labels, arXiv:2406.17633. Preprint, arXiv, 25 June 2024. 10.48550/arXiv.2406.17633.

Przybyla L, Gilbert LA. A new era in functional genomics screens. Nat Rev Genet 2022;23(2):89– 103. 10.1038/s41576-021-00409-w.

Rahit Kmth, Avramovic V, Chong JX et al. GPAD: a natural language processing-based application to extract the gene-disease association discovery information from OMIM. BMC Bioinformatics 2024;25(1):84. 10.1186/s12859-024-05693-x.

Rai P, Jain A, Kumar S et al. Literature mining discerns latent disease–gene relationships. Bioinformatics 2024;40(4):btae185. 10.1093/bioinformatics/btae185.

Ravikumar KE, Rastegar-Mojarad M, Liu H. BELMiner: adapting a rule-based relation extraction system to extract biological expression language statements from bio-medical literature evidence sentences. Database 2017;2017:baw156. 10.1093/database/baw156.

Schipper M, Leeuw CA de, Maciel Bapc et al. Prioritizing effector genes at trait-associated loci using multimodal evidence. Nat Genet 2025;57(2):323–33. 10.1038/s41588-025-02084-7.

Schmidt MB, Chen EH, Lynch SE. A review of the effects of insulin-like growth factor and platelet derived growth factor on in vivo cartilage healing and repair. Osteoarthritis Cartilage 2006;14(5):403–12. 10.1016/j.joca.2005.10.011.

Sener O, Savarese S. Active Learning for Convolutional Neural Networks: A Core-Set Approach, arXiv:1708.00489. Preprint, arXiv, 1 June 2018. 10.48550/arXiv.1708.00489.

Shelmanov A, Liventsev V, Kireev D et al. Active Learning with Deep Pre-trained Models for Sequence Tagging of Clinical and Biomedical Texts. 2019 IEEE Int Conf Bioinforma Biomed BIBM Nov. 2019:482–9. 10.1109/BIBM47256.2019.8983157.

Shi Z, Li M, Zhou H. A Weighted BioBERT and Active Learning based Biomedical Named Entity Recognition Method with Small Amount of Labeled Data. 2023 Int Conf Intell Commun Netw ICN Nov. 2023:244–8. 10.1109/ICN60549.2023.10426620.

Soul J, Young DA. Machine learning based prioritisation of genes associated with osteoarthritis joint damage in animals. Osteoarthritis Cartilage 2026;34(3):475–83. 10.1016/j.joca.2026.01.002.

Suvalov H, Laur S, Kolde R. Information Extraction from Medical Texts with BERT Using Human-in-the-Loop Labeling. Stud Health Technol Inform 2023;302:831–2. 10.3233/SHTI230281.

Team G, Mesnard T, Hardin C et al. Gemma: Open Models Based on Gemini Research and Technology, arXiv:2403.08295. Preprint, arXiv, 16 Apr. 2024. 10.48550/arXiv.2403.08295.

The Gene Ontology Consortium. The Gene Ontology knowledgebase in 2026. Nucleic Acids Res 2026;54(D1):D1779–92. 10.1093/nar/gkaf1292.

Tomanek K, Hahn U. Reducing class imbalance during active learning for named entity annotation. Proc Fifth Int Conf Knowl Capture (New York, NY, USA) K-CAP ‘09, 1 Sept. 2009:105–12. 10.1145/1597735.1597754.

Touvron H, Lavril T, Izacard G et al. LLaMA: Open and Efficient Foundation Language Models, arXiv:2302.13971. Preprint, arXiv, 27 Feb. 2023. 10.48550/arXiv.2302.13971.

Van Auken K, Schaeffer ML, McQuilton P et al. BC4GO: a full-text corpus for the BioCreative IV GO task. Database 2014;2014:bau074. 10.1093/database/bau074.

Vollmar M, Tirunagari S, Harrus D et al. Dataset from a human-in-the-loop approach to identify functionally important protein residues from literature. Sci Data 2024;11(1):1032. 10.1038/s41597-024-03841-9.

Wehling N, Palmer GD, Pilapil C et al. Interleukin-1β and tumor necrosis factor α inhibit chondrogenesis by human mesenchymal stem cells through NF-κB–dependent pathways. Arthritis Rheum 2009;60(3):801–12. 10.1002/art.24352.

Wei CH, Luo L, Islamaj R et al. GNorm2: an improved gene name recognition and normalization system. Bioinformatics 2023;39(10):btad599. 10.1093/bioinformatics/btad599.

Welsh BL, Sikder P. Advancements in Cartilage Tissue Engineering: A Focused Review. J Biomed Mater Res B Appl Biomater 2025;113(1):e35520. 10.1002/jbm.b.35520.

Wu Y, Xu J, Li J et al. Deep graph convolutional network-based multi-omics integration for cancer driver gene identification. Brief Bioinform 2025;26(4):bbaf364. 10.1093/bib/bbaf364.

Xing W, Qi J, Yuan X et al. A gene–phenotype relationship extraction pipeline from the biomedical literature using a representation learning approach. Bioinformatics 2018;34(13):i386– 94. 10.1093/bioinformatics/bty263.

Yadav S, Ramesh S, Saha S et al. Relation Extraction From Biomedical and Clinical Text: Unified Multitask Learning Framework. IEEE/ACM Trans Comput Biol Bioinform 2022;19(2):1105–16. 10.1109/TCBB.2020.3020016.

Yang A, Li A, Yang B et al. Qwen3 Technical Report, arXiv:2505.09388. Preprint, arXiv, 14 May 2025. 10.48550/arXiv.2505.09388.

Yasunaga M, Leskovec J, Liang P. LinkBERT: Pretraining Language Models with Document Links. Proc 60th Annu Meet Assoc Comput Linguist Vol 1 Long Pap 2022:8003–16. 10.18653/v1/2022.acl-long.551.

Yu E, Larivie re R, Thomas RA et al. Machine learning nominates the inositol pathway and novel genes in Parkinson’s disease. Brain 2024;147(3):887–99. 10.1093/brain/awad345.

Zhao XM, Wang Y, Chen L et al. Gene function prediction using labeled and unlabeled data. BMC Bioinformatics 2008;9(1):57. 10.1186/1471-2105-9-57.

Zhao Y, Goto S. Can Frontier LLMs Replace Annotators in Biomedical Text Mining? Analyzing Challenges and Exploring Solutions, arXiv:2503.03261. Preprint, arXiv, 17 May 2025. 10.48550/arXiv.2503.03261.

Zhou D, He Y, Kwoh CK. Extracting Protein-Protein Interactions from the Literature Using the Hidden Vector State Model. In: Alexandrov VN, Albada GD van, Sloot PMA et al. (eds), Computational Science – ICCS 2006. Berlin, Heidelberg: Springer, 2006, 718–25. 10.1007/11758525_97.

