## Supplementary Material for "A low-annotation-budget PubMedBERT classifier for chondrogenesis regulator discovery via active learning"

### Supplementary Tables

***Table S1. Filter terms applied to retrieved sentences that contain at least one term from the sentences extracted from PubMed, grouped by category.***

| Category | Terms |
|---|---|
| **Cell and tissue identity** | cartilage, chondrocyte, chondrocytic, chondrogenic |
| **Developmental process** | chondrogenesis, development, differentiate, differentiation, growth |
| **Cell state and expansion** | proliferation, hypertrophy, hypertrophic |
| **Repair and regeneration** | repair, regeneration, regenerate, regenerates |
| **Evidence type** | markers |

***Table S2. Entity counts at each stage of the text-processing pipeline for the three datasets, along with the specific query used in this study: abstracts retrieved, sentences extracted, sentences retained after term filtering, and gene mentions identified.***

| Data Set | PubMed Extraction Query | Number of articles extracted | Number of sentences after splitting | Number of sentences after filtering | Number of gene mentions |
|---|---|---|---|---|---|
| **Training-validation-LLM Dev** | '((chondrogenesis OR chondrocyte differentiation) AND "English"[Language]) NOT "Review"[Publication Type] AND ("1000/01/01"[dp] : "2025/01/31"[dp])' | 17,316 | 185,704 | 39,180 | 85,004 |
| **Held out** | '((chondrogenesis OR chondrocyte differentiation) AND "English"[Language]) NOT "Review"[Publication Type] AND ("2025/06/01"[dp] : "2025/12/31"[dp])' | 395 | 4,553 | 835 | 1,827 |
| **Application** | '((chondrogenesis OR chondrocyte differentiation) AND "English"[Language]) NOT "Review"[Publication Type] AND (“2006/01/01"[dp] : "2026/02/28"[dp])' | 13,877 | 152,431 | 31,325 | 67,111 |

*Table S3. Number of sentences selected at each active learning iteration using different sampling strategies.*

| Type of Sampling | Core | Iter 1 | Iter 2 | Iter 3 | Iter 4 | Iter 5 |
|---|---|---|---|---|---|---|
| **Diversity** | 100 | 100 | 50 | 50 | 20 | 20 |
| **Uncertainty** | 0 | | 50 | 50 | 70 | 70 |
| **Random** | 0 | 0 | 0 | 0 | 10 | 10 |

*Table S4. Python environment specifications for the three model families evaluated. The active learning environment was used for PubMedBERT training and inference, the Llama environment for the quantised Llama-3.1-70B, and the Qwen environment for all Qwen3 models. Separate environments were required because of incompatible dependency requirements across model families.*

| Package | AL environment | Llama environment | Qwen Environment |
|---|---|---|---|
| **Torch** | 2.10.0 | 2.7.0 | 2.13.0 |
| **Transformers** | 4.51.0 | 4.52.3 | 5.5.0 |
| **huggingface** | 0.36.2 | 1.23.0 | 0.32 |

*Table S5. Hyperparameter search space for fine-tuning the PubMedBERT classifier. Each configuration was evaluated by stratified 5-fold cross-validation*

| Parameter | Options |
|---|---|
| **Learning rate** | 1e-5, 4e-5, 7e-5, 1e-4 |
| **Number of epochs** | 2, 3, 4 |
| **Weight decay** | 0.01, 0.001 |
| **Batch size** | 8, 16, 32, 48 |
| **Warmup steps** | 0, 0.05, 0.07, 0.1 |
| **Class-balanced loss function** | True, False |

*Table S6. Performance and computational resources comparison of all evaluated models on the held-out dataset*

| Model | Parameters | Prompting | Tuning Variant | Training Time (s) | Average Gene Inference Time (ms) | Projected corpus runtime (67111 genes) | Precision | Recall | F1 | AUC-ROC |
|---|---|---|---|---|---|---|---|---|---|---|
| **Logistic Regression** | - | - | - | 36.86 | 0.003 | 0.2s | 0.35 [0.09, 0.64] | 0.09 [0.02, 0.17] | 0.14 [0.03, 0.26] | 0.69 [0.61, 0.76] |
| **PubMedBERT** | 109M | - | - | 88.58 | 4.1 | 4.6min | 0.69 [0.56, 0.82] | 0.66 [0.53, 0.78] | 0.67 [0.56, | 0.93 [0.89, |

| | | | | | | | | | |
|---|---|---|---|---|---|---|---|---|---|
| | | | | | | | | 0.77] | 0.96] |
| **Qwen3** | 4B | Few-shot | Instruction | - | 890 | 16.59h | 0.43 [0.34, 0.52] | 0.68 [0.57, 0.8] | 0.52 [0.43, 0.61] | 0.86 [0.8, 0.9] |
| **Qwen3** | 4B | Zero-shot | Instruction | - | 560 | 10.44h | 0.45 [0.35, 0.56] | 0.66 [0.55, 0.76] | 0.53 [0.44, 0.62] | 0.86 [0.81, 0.9] |
| **Qwen3** | 30B | Few-shot | Instruction | - | 1,098 | 20.47h | 0.42 [0.33, 0.52] | 0.85 [0.76, 0.94] | 0.57 [0.47, 0.65] | 0.91 [0.88, 0.94] |
| **Qwen3** | 30B | Zero-shot | Instruction | - | 458 | 8.54h | 0.47 [0.37, 0.57] | 0.78 [0.66, 0.89] | 0.59 [0.49, 0.68] | 0.91 [0.87, 0.94] |
| **Llama3.1** | 70B | Few-shot | Instruction | - | 1,822 | 1.42d | 0.43 [0.35, 0.52] | 0.93 [0.85, 0.99] | 0.59 [0.5, 0.68] | 0.94 [0.91, 0.96] |
| **Llama3.1** | 70B | Zero-shot | Instruction | - | 1,277 | 23.81h | 0.40 [0.31, 0.49] | 0.91 [0.84, 0.98] | 0.56 [0.46, 0.64] | 0.94 [0.91, 0.96] |
| **Qwen3** | 4B | Few-shot | Reasoning | - | 118,746 | 92.24d | 0.64 [0.52, 0.76] | 0.74 [0.63, 0.85] | 0.69 [0.59, 0.78] | 0.85 [0.79, 0.91] |
| **Qwen3** | 4B | Zero-shot | Reasoning | - | 91,510 | 71.08d | 0.67 [0.55, 0.78] | 0.67 [0.55, 0.78] | 0.67 [0.57, 0.76] | 0.86 [0.81, 0.91] |
| **Qwen3** | 30B | Few-shot | Reasoning | - | 137,321 | 106.66 d | 0.68 [0.57, 0.78] | 0.78 [0.67, 0.88] | 0.73 [0.63, 0.81] | 0.94 [0.91, 0.97] |
| **Qwen3** | 30B | Zero-shot | Reasoning | - | 105,520 | 81.96d | 0.74 [0.63, 0.85] | 0.74 [0.62, 0.86] | 0.74 [0.64, 0.83] | 0.93 [0.89, 0.97] |

### Supplementary Notes

***Note S1. Annotation Guidelines***

#### General rule

A chondrogenesis regulator is a gene whose activity directly modulates the frequency, rate, or extent of chondrogenesis. We should label a gene mention as a *Driver* if there is explicit, non-speculative evidence that the gene directly regulates chondrogenesis, and as a *Non-Driver* if such evidence is absent.

#### Rule Refinements

| Rule | Description | Example Case | Gene to classify | Label |
|---|---|---|---|---|
| ***Clear Gene-Chondrogenesis Relationship*** | The relationship between the gene and chondrogenesis must be explicitly stated in the text. Do not rely on external knowledge or inference to determine relevance | In addition, caMsx2 overexpression induced Ihh (Indian hedgehog) expression in mouse primary chondrocytes | caMsx2 | Non-Driver |
| ***Direct Relation to Chondrogenesis Only*** | Only genes that are directly involved in chondrogenesis (or chondrogenesis potential) should be labelled as related. Associations with general proliferation indicators should not be considered unless it is specified that it is chondrogenesis related | Over-expression of Aire induced the early stages of chondrocyte differentiation by facilitating expression of Bmp2 | Aire | Driver |
| ***Handling of Model Verbs*** | Genes that use modal verbs such as "may", "might" or "could" are considered speculative and should not be classified as evidence of chondrogenesis | Differential mRNA expression of TGF-beta1 was found during all passages, which suggests that this growth factor might be involved in chondrocyte. | TGF-beta1 | Non-Driver |
| ***Handling of negative evidence*** | Do NOT consider negative evidence. If a gene "supports another process" and does not influence chondrogenesis directly, it is not counted as being involved in chondrogenesis | Knockdown of FOXA2 promoted osteogenic differentiation of bone-marrow-derived mesenchymal stem cells while not altering chondrogenic potential | FOXA2 | Non-Driver |
| ***Pathways and axis*** | Genes that are part of chondrogenic pathways or axes should not be classified as drivers when mentioned as intermediates or | Cytl1 exerted its chondrogenic effect via stimulation of Sox9 transcriptional activity | Cytl1 | Driver |
| | | | Sox9 | Non-Driver |

| | | | | |
|---|---|---|---|---|
| | downstream markers. In such cases, they serve as evidence (phenotypic or biomarker proof) that another gene acts as a regulator of chondrogenesis | | | |
| ***Exclusion of Hypertrophy and Osteogenesis*** | Genes related to hypertrophy (also referred to as "terminal chondrogenesis" or "terminal maturation" or reduced osteogenesis should not be labeled as related to chondrogenesis. | FOXC1 and FOXC2 regulate growth plate chondrocyte maturation towards hypertrophy in the embryonic mouse limb skeleton. | FOXC1 | Non-Driver |
| ***Exclusion of Cell Proliferation, Re-differentiation and Repair*** | Genes related to only cell proliferation or cartilage repair or re-differentiation should not be labeled as related to chondrogenesis | We found that silencing of miR-221 strongly enhanced in vivo cartilage repair compared to the control conditions | miR-221 | Non-Driver |
| ***Cross-Species Consideration*** | Information from all species is valid and should be taken into account when labeling | Importantly, TET1 inhibition in vivo in late stages of a mouse model of OA led to increased cartilage regeneration | TET1 | Non-Driver |
| ***Cell types Consideration*** | We will consider all types of cells-lines for the search of chondrogenesis drivers | In Conclusion, our data implies that miR-140 is a potent chondrogenic differentiation inducer for iPSCs and also, we have showed increasing chondrogenic differentiation by using overexpression of miR-140 and TGFb3 | TGFb3 | Non-Driver |
| ***Tri-lineage cells Consideration*** | Any sentence mentioning "tri-lineage cells" should be treated as indicating chondrocytes, osteoblasts and adipocytes, since our data acquirement was focused on chondrogenesis | We have shown that overexpression of Sox11 in rMSCs by lentivirus-mediated gene transfer leads to enhanced tri-lineage differentiation and accelerated bone formation in fracture model of rats | Sox11 | Driver |
| ***All biomolecules Considered*** | We will be considering a less strict definition of gene. This means | Knockdown of miR-145 promoted chondrogenesis and inhibited hypertrophy differentiation in RMCs | miR-145 | Driver |

| | | | | |
|---|---|---|---|---|
| | that protein-coding and non-coding genes will be considered, in addition to molecules such as microRNAs | | | |
| ***Bio-markers relations*** | We will be considering drivers related to the expression of chondrogenesis biomarkers such as ACAN, SOX9, etc only if there is another specification in the text that they are specific chondrogenesis markers. We will not consider the genes drivers if the already known biomarkers are by themself. If in the text only biomarkers are shown as proof, the biomarker proof box needs to be selected when labelling the text | Furthermore, Arid5a physically interacted with Sox9 in nuclei and up-regulated the chondrocyte-specific action of Sox9 | Arid5a | Driver |
| | | Furthermore, Arid5a physically interacted with Sox9 in nuclei and up-regulated the action of Sox9 | Arid5a | Non-Driver |
| ***Mentioning of cartilage associated diseases*** | We will not consider markers of diseases related to chondrogenesis. Only because the disease is associated to chondrogenesis, we will not take it as a driver of this process | Inhibiting the expression of CDKN1A can significantly suppress the differentiation of OA chondrocytes | CDKN1A | Non-Driver |
| ***Functional Role vs Marker Role*** | Only label genes that are described as having a direct functional or regulatory role in chondrogenesis. If a gene is mentioned only as a marker or indicator of chondrogenesis, do not label it as directly involved | Genome-wide mRNA expression analysis of the midpalatal suture tissue revealed that MLL4 is essential for the timely expression of major cartilage development genes, such as Col2a1 and Acan, at birth | MLL4 | Driver |
| | | | Col2a1 | Non-Driver |

| | | | | |
|---|---|---|---|---|
| ***Growth Plate Mention*** | If the mention refers to the growth plate, it must be explicitly stated that the process involves chondrocytes | Sox9 is involved in the process of endochondral ossification within the growth plate | Sox9 | Non-Driver |

##### *Note S2. System prompt*

<role>

You are a molecular biology assistant that works in chondrogenesis.

</role>

<task>

Classify whether a gene is a regulator of chondrogenesis.

</task>

<context>

<definition_chondrogenesis>

Chondrogenesis is the biological process through which cartilage tissue, known as chondrocytes, is formed and developed.

</definition_chondrogenesis>

<definition_regulator>

A chondrogenesis regulator is a gene whose activity directly modulates the frequency, rate, or extent of chondrogenesis.

</definition_regulator>

</context>

<rules_for_classification>

<general_decision_rules>

<rule>Output "Yes" only if there is explicit, non-speculative evidence that the gene directly regulates chondrogenesis.</rule>

<rule>Output "No" if explicit, non-speculative evidence that the gene directly regulates chondrogenesis is absent.</rule>

</general_decision_rules>

<evidence_requirements>

<rule>The regulatory relationship between the gene and chondrogenesis must be explicitly stated in the text.</rule>

<rule>Acceptable evidence includes direct experimental or mechanistic statements (e.g., promotes, inhibits, regulates, is required for chondrogenesis).</rule>

<rule>Acceptable evidence also includes phenotypic evidence demonstrating that manipulation of the gene alters chondrogenic outcomes (e.g., cartilage formation).</rule>

<rule>Acceptable evidence also includes changes in chondrogenesis molecular markers when the gene is experimentally manipulated.</rule>

<rule>Acceptable evidence also includes the activation or inhibition of pathways or signaling cascades that lead to or are required for chondrogenesis.</rule>

<rule>Acceptable evidence also includes affirmative statements attributing a role in chondrogenesis to the target gene, even when established or background knowledge rather than the result of a direct experiment.</rule>

<rule>Speculative language (modal verbs) does not qualify as evidence when the sentence is framing a hypothesis or a future possibility.</rule>

<rule>Use only the information explicitly stated in the provided text.</rule>

<rule>If the sentence does not contain enough evidence, the gene is not a regulator.</rule>

<rule>Being activated during chondrogenesis does not imply regulation of chondrogenesis.</rule>

</evidence_requirements>

<process_constraints>

<rule>Regulation must involve chondrogenesis, cartilage development, chondrocyte maturation, chondrocyte differentiation or chondrogenic potential of chondrocytes.</rule>

<rule>Genes involved only in cell proliferation, cell condensation, chondrocyte proliferation, hypertrophy, terminal maturation, terminal differentiation, cartilage repair, cartilage regeneration, or cartilage redifferentiation are not regulators.</rule>

<rule>Genes acting only as downstream markers or as intermediate effectors in a signaling cascade are not regulators, only the upstream initiating gene driving the cascade is considered a regulator.</rule>

<rule>Genes used as markers of chondrogenesis (e.g., to confirm or assess chondrogenic differentiation) are not regulators, even if they are expressed during the process.</rule>

<rule>Genes associated only with cartilage-related diseases are not regulators.</rule>

<rule>Growth plate–related genes are regulators only if chondrocyte involvement is explicitly stated.</rule>

</process_constraints>

<biological_context>

<rule>Genes from all species are valid.</rule>

<rule>Genes from all cell lines are valid.</rule>

<rule>Mentions of tri-lineage differentiation imply chondrocytes, osteoblasts, and adipocytes.</rule>

</biological_context>

<target_specificity>

<rule>Only consider evidence related to the gene marked as [TARGET][GENE][/TARGET].</rule>

<rule>Only consider statements that explicitly describe the role of the [TARGET][GENE][/TARGET].</rule>

<rule>Ignore information about other genes mentioned in the sentence.</rule>

<rule>If the target gene is merely regulated by another gene, this does not imply that it regulates chondrogenesis unless explicitly stated.</rule>

<rule>Each masked gene must be treated as an independent entity, do not assume that different masked genes in different part of the text are the same ones.</rule>

</target_specificity>

</rules_for_classification>

<input>

You will receive sentences containing one or more genes marked with [GENE].

Only one gene will be marked as [TARGET][GENE][/TARGET], which is the gene to classify.

</input>

<masked_entity_handling>

<rule>All genes are masked and have no known identity.</rule>

<rule>Do not infer the target gene identity from contextual information</rule>

<rule>Classification must be based solely on the explicit role described in the sentence.</rule>

<rule>Do not use prior or external knowledge about the target gene's known biological functions when making the classification decision.</rule>

</masked_entity_handling>

<output>

Output must consist of a single word only: "Yes" or "No".

Output "Yes" if the target gene is a regulator of chondrogenesis.

Output "No" otherwise.

</output>

***Note S3. Few Shot User Prompt***

**## Task**

You will be provided with text from research articles containing one or more genes marked as [GENE]. One gene is marked as [TARGET][GENE][/TARGET] and is referred to as the target gene.

Your task is to determine whether the target gene is a regulator of chondrogenesis based only on the provided text.

**## Output**

Output must be a single word: Yes or No.

Do not provide explanations or additional text. Do not restate the sentence. Do not mention any genes by name.

**## Examples**

1. [TARGET][GENE][/TARGET] cannot be substituted by [GENE], but dexamethasone concentration can be decreased to 10(-12) M without chondrogenesis being impaired. // Yes

2. Our results suggest that [TARGET][GENE][/TARGET] acts as a positive regulator for [GENE] expression and may cooperate with [GENE] together to control [GENE] expression and promote the proliferation and maturation of chondrocytes. // Yes

3. The other results demonstrated that higher the ATDC5 ratios and longer the culture duration, greater the expression of cartilage-specific genes (including [GENE] and [TARGET][GENE][/TARGET]) and more the synthesized cartilaginous extracellular matrix. // No

4. [GENE] ([GENE]) inhibitors are compounds that can induce the [TARGET][GENE][/TARGET] signaling pathway, which is involved in chondrogenesis and cartilage development. // Yes

5. Although [TARGET][GENE][/TARGET] is thought to be specifically required for chondrogenesis, human fetuses with the mutation of [GENE] also display bony skull defects where chondrocytes are usually not present. // Yes

6. Although [GENE] is thought to be specifically required for chondrogenesis, human fetuses with the mutation of [TARGET][GENE][/TARGET] also display bony skull defects where chondrocytes are usually not present. // No

7. [GENE] ([TARGET][GENE][/TARGET]) is dysregulated in some cancers and may regulate cell proliferation in specific contexts. // No

8. [TARGET][GENE][/TARGET] reduced the chondrogenic potential of these cells via the upregulation of [GENE] protein and enhanced [GENE] protein and [GENE] mRNA levels. // Yes

9. RESULTS: Downregulation of [TARGET][GENE][/TARGET] expression in [GENE] ([GENE])-stimulated chondrocyte pellet cultures led to chondrocyte terminal differentiation characterized by poor production of cartilage extracellular matrix and altered expression of genes and proteins involved in cartilage homeostasis, including [GENE], [GENE], [GENE], [GENE], [GENE], [GENE], and [GENE]. // No

10. [GENE] ([TARGET][GENE][/TARGET]) plays an essential role in chondrocyte maturation. // Yes

11. [TARGET][GENE][/TARGET] and [GENE] ([GENE])-[GENE] signaling play important roles in osteoblast and chondrocyte differentiation. // Yes

**## Text to classify**

{sentence_to_classify}

***Note S4. Zero Shot User prompt***

**## Task**

You will be provided with text from research articles containing one or more genes marked as [GENE]. One gene is marked as [TARGET][GENE][/TARGET] and is referred to as the target gene.

Your task is to determine whether the target gene is a regulator of chondrogenesis based only on the provided text.

**## Output**

Output must be a single word: Yes or No.

Do not provide explanations or additional text. Do not restate the sentence. Do not mention any genes by name.

**## Text to classify**

{sentence_to_classify}

### Supplementary Figures

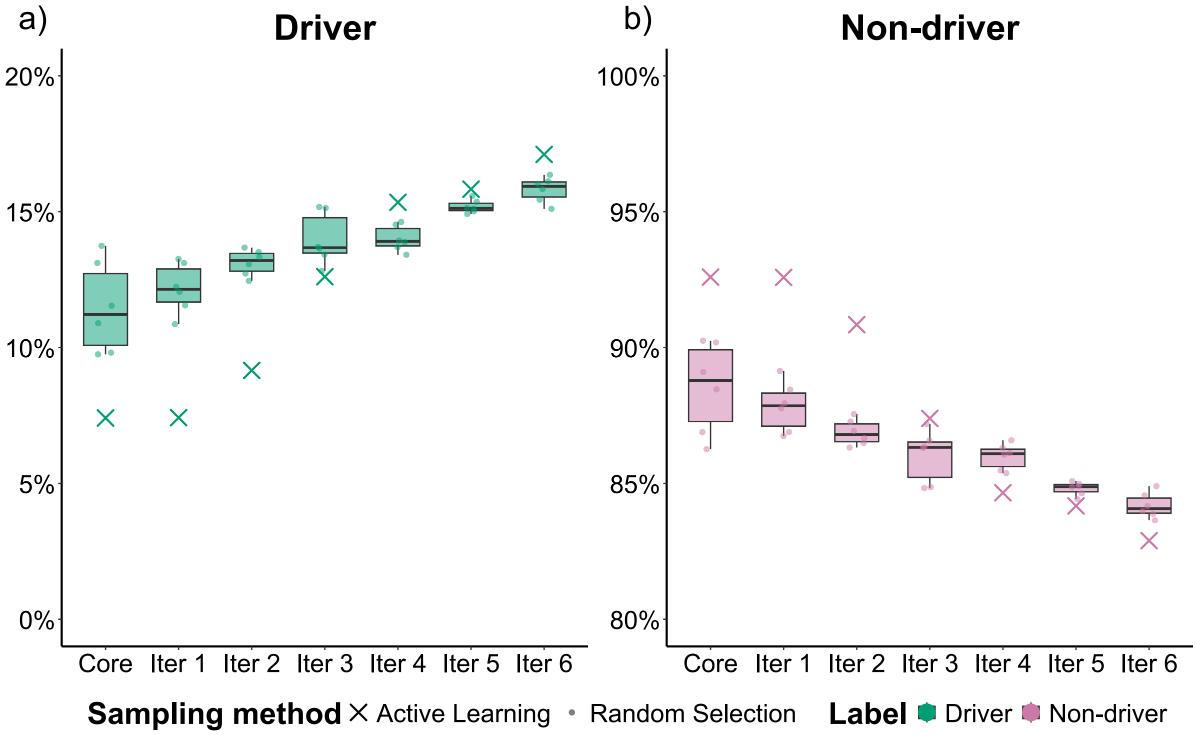


***Figure S1. Composition of the training dataset for random sampling and active learning models.*** *Distribution of Driver and Non-driver labels in the sampled annotation sets across the same iterations. Boxplots show the distribution across seeds for the random-sampling baseline; crosses show the composition of the active-learning training set at each iteration.*

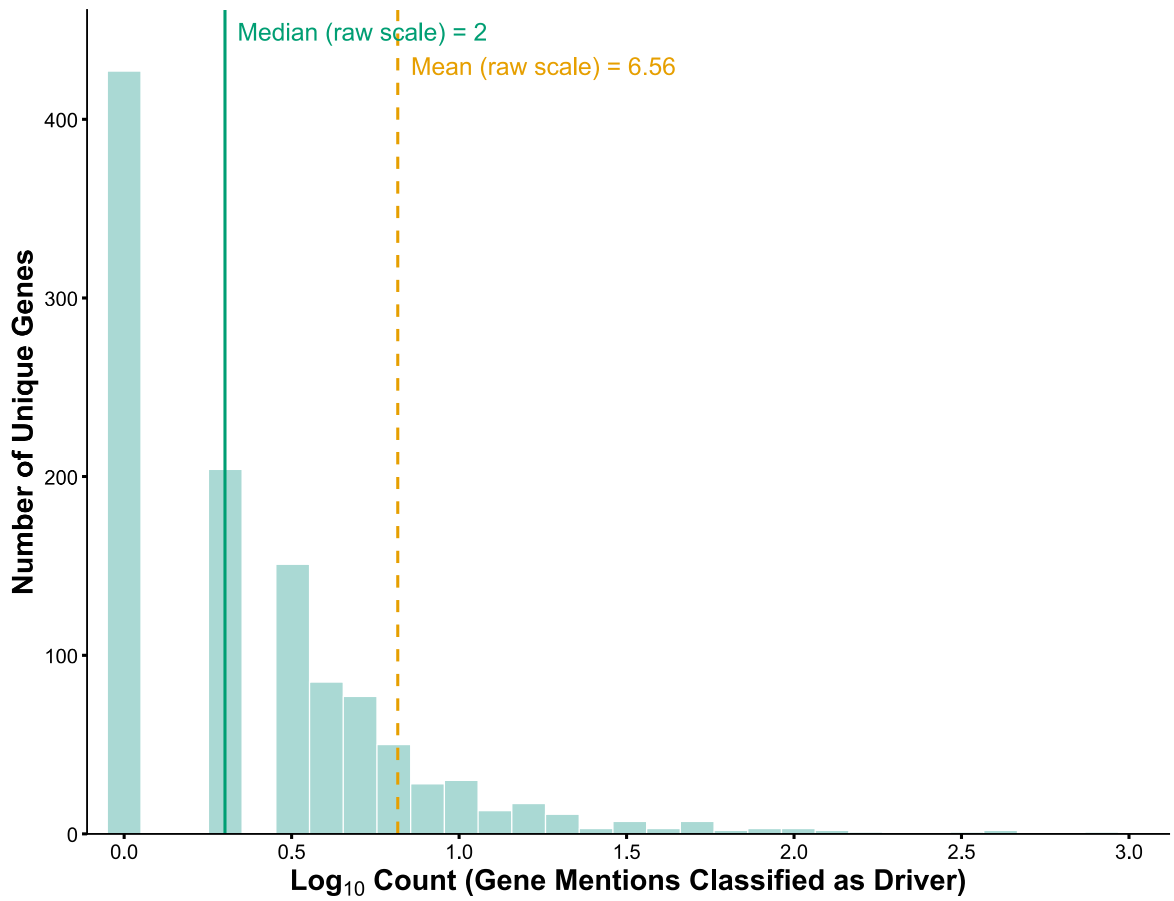


***Figure S2. Distribution of gene mention frequencies classified as chondrogenesis drivers for the normalised genes.*** *Histogram displaying the log10-transformed driver-classified gene mention counts of normalised genes. The solid green vertical line represents the data median in raw units (counts), and the orange dashed line represents the data mean.*

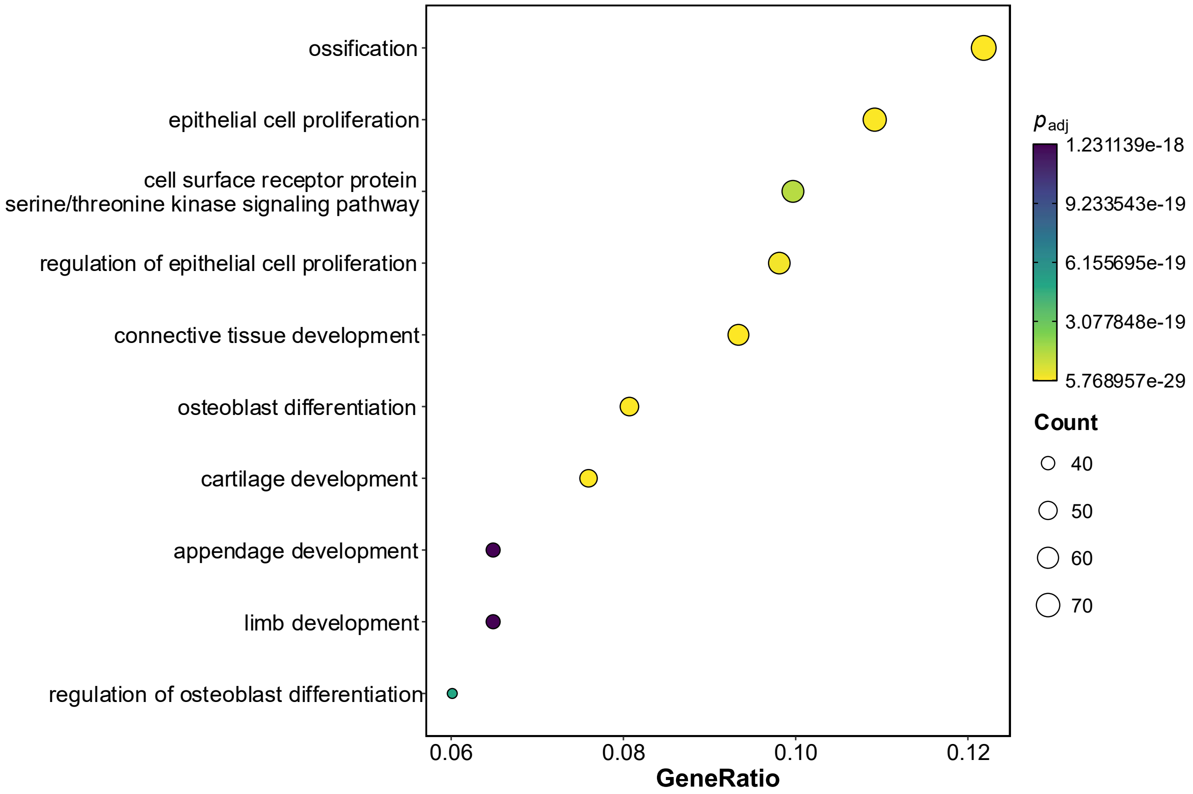


***Figure S3. Gene Ontology Biological Process enrichment analysis of gene mentions classified as drivers of chondrogenesis.*** *Dot plot of the top 10 significantly enriched Gene Ontology Biological Process terms from over-representation analysis (ORA). Dot size reflects gene count per term, and colour reflects the Benjamini-Hochberg-adjusted p-value.*
